# Purification of Cross-linked RNA-Protein Complexes by Phenol-Toluol Extraction

**DOI:** 10.1101/333385

**Authors:** Erika C Urdaneta, Carlos H Vieira-Vieira, Timon Hick, Hans-Herrmann Wessels, Davide Figini, Rebecca Moschall, Jan Medenbach, Uwe Ohler, Sander Granneman, Matthias Selbach, Benedikt M Beckmann

## Abstract

Recent methodological advances allowed the identification of an increasing number of RNA-binding proteins (RBPs) and their RNA-binding sites. Most of those methods rely, however, on capturing proteins associated to polyadenylated RNAs which neglects RBPs bound to non-adenylate RNA classes (tRNA, rRNA, pre-mRNA) as well as the vast majority of species that lack poly-A tails in their mRNAs (including all archea and bacteria). To overcome these limitations, we have developed a novel protocol, Phenol Toluol extraction (PTex), that does not rely on a specific RNA sequence or motif for isolation of cross-linked ribonucleoproteins (RNPs), but rather purifies them based entirely on their physicochemical properties. PTex captures RBPs that bind to RNA as short as 30 nt, RNPs directly from animal tissue and can be used to simplify complex workflows such as PAR-CLIP. Finally, we provide a first global RNA-bound proteome of human HEK293 cells and *Salmonella* Typhimurium as a bacterial species.

---

RNA binding proteins are key factors in the post transcriptional regulation of gene expression. Spurred by recent technological advances such as RNA interactome capture (RIC) (1–3), the number of RBPs has been greatly increased. A powerful tool to study ribonucleoproteins (RNPs) is UV cross-linking: irradiation of cells with short wavelength UV light results in covalent cross-links of proteins in direct contact with the RNA (Fig. 1a) (4–6). Exploiting the stability of cross-linked RNPs, new methods have been developed to identify and analyse RNPs: i) RNA interactome capture in which poly-A RNA and its bound proteins are first selected by oligo-dT beads and co-purified proteins subsequently identified by mass spectrometry. This lead to the discovery of hundreds of hitherto unknown RBPs (7, 8). ii) CLIP (cross-linking and immunoprecipitation) and similar methods in which, after UV cross-linking, individual RBPs are immunoprecipitated, and co-precipitated transcripts are identified by RNA-Seq, yielding high resolution data on the RNA binding sites of the RBPs of interest (9–15).

**Fig. 1.**
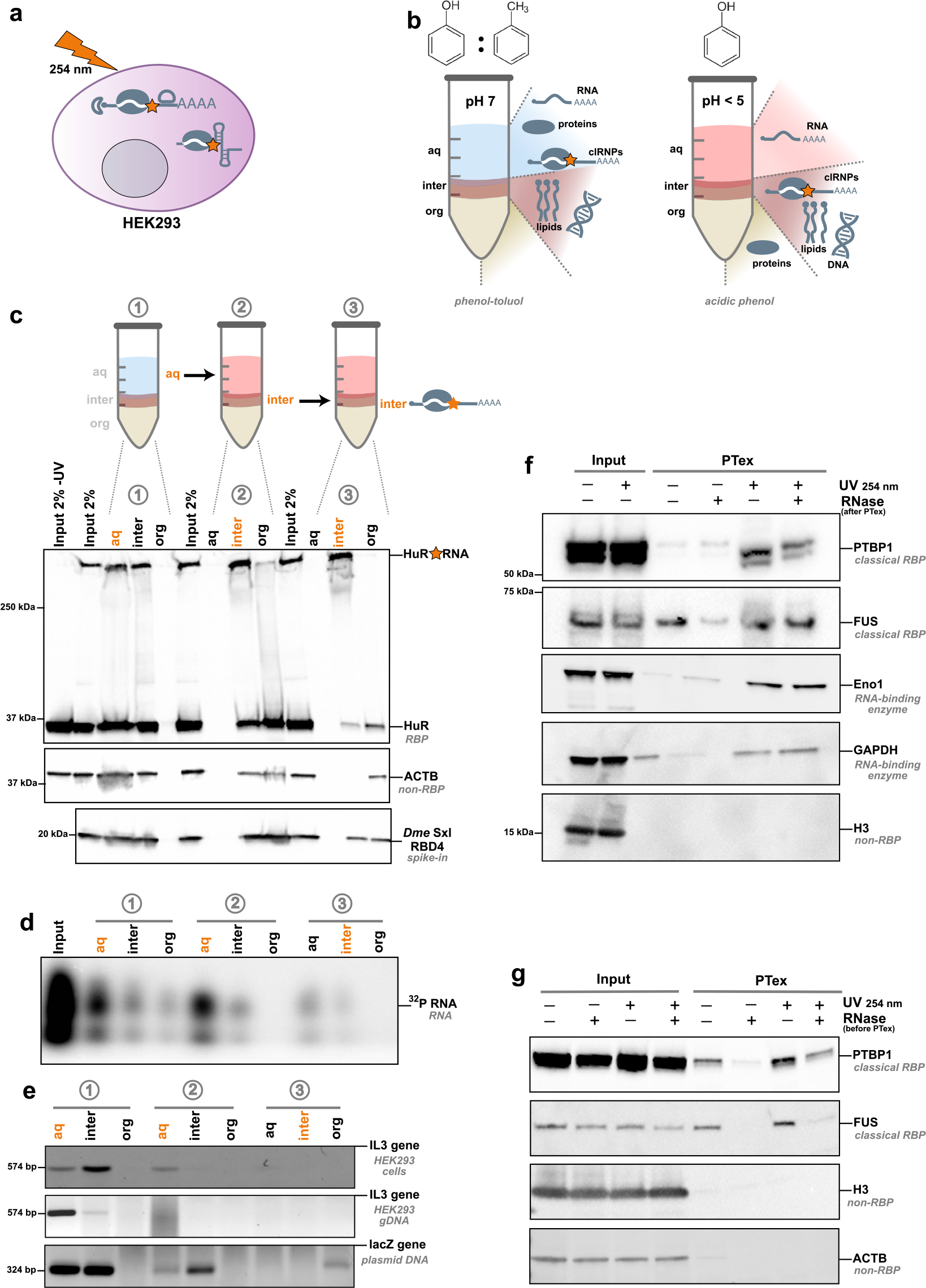
PTex is a fast method to purify cross-linked RNPs. a) *In vivo* cross-linking of HEK293 cells using UV light at 254 nm wavelength results in covalent bonds between RNA and proteins in direct contact. Cross-linked RNPs are indicated by an orange star. b) Schematic of the separation principle of biphasic organic extractions used in PTex. Left panel: Phenol-Toluol (50:50) and neutral pH results in an accumulation of proteins and RNA in the upper aqueous phase (aq) while DNA and lipids are retained at the interphase (inter). Right panel: under acidic phenol and chaotropic conditions, non-cross-linked RNA accumulates in the aqueous phase (aq), non-cross-linked proteins in the lower organic phase (org) and cross-linked RNPs (clRNPs) are enriched at the interphase (inter). c) Step-by-step analysis of proteins in 9 intermediary steps of the PTex protocol (3 extractions with 3 phases each). Western blot against HuR (ELAVL1, 35 kDa) demonstrates that UV-cross-linking-stabilised HuR-RNA complexes (upper edge/gel pocket of the blot; see Supplementary Figure S1; cross-linking is denoted with an orange star) are largely enriched after PTex (step 3 interphase). Abundant cellular non-RNA-binders such as beta-actin (ACTB) are efficiently removed by PTex. A purified fly protein (Sxl RBD4) served as spike-in as 100% non-cross-linked RBP; ~99% of the free protein is removed by PTex. d) Step-by-step analysis of RNA. 5‘-end radioactive labeled RNA was subjected to PTex *in vitro*. Like proteins, non-cross-linked RNA is efficiently depleted in the PTex fraction (step 3 interphase). e) Testing for DNA by PCR with specific primers against exon 5 of the interleukin 3 (IL3) gene demonstrates efficient removal of genomic DNA after either full HEK293 cells (upper panel) or pre-purified genomic DNA (middle panel) were subjected to PTex. A PCR product derived from linear pUC19 DNA (lower panel) is also removed. f) Enrichment of known RBPs by PTex tested by western-blot against PTBP1 and FUS, or against “non-classical” RNA-binding enzymes Eno1 and GAPDH. Note that RNaseA treatment was performed after PTex as it removes partially shifted bands (smear) for some RBPs. g) PTex enriches for cross-linked RBPs. RNase treatment before PTex strongly reduces recovery of known RNA-binders (PTBP1, FUS). Non-RBP controls Histone H3 and actin (ACTB) are efficiently depleted by PTex (f,g). For full gels/blots see Supplementary Figures S2-S8.

As RNA interactome capture relies on the purification of cross-linked RNPs based on hybridisation of oligo-dT beads to oligo-A sequences typically found in eukaryotic messenger RNAs, RBPs that exclusively associate with non-adenylate RNA species such as e.g. rRNA, tRNAs, snRNAs, histone mRNAs, or numerous lncRNAs cannot be identified.

The same limitations apply to mRNA from bacteria and archaea that lack poly-A tails in general. Recently, RNA interactome using click chemistry (RICK (16) and CARIC (17)) has been introduced in which labeled RNA along with UV-cross-linked interacting proteins was purified in a poly-A-independent fashion. However, the method requires efficient *in vivo* labeling of RNA, limiting its application to suitable (cell culture) systems. Consequentially, no RNA-bound proteomes of prokaryotes have been determined by biochemical means to date.

A commonly used protocol to purify RNA from whole cell lysates is the single step method (18), also marketed as “Trizol”. First, chaotropic conditions and ionic detergents are employed to denature cellular components, followed by a biphasic extraction using the organic compound phenol. During this treatment, nucleic acids are specifically enriched in the aqueous phase. Furthermore, the pH during extraction allows to control if DNA and RNA (neutral pH) or only RNA (acidic pH) accumulate in the aqueous phase (acidic conditions shown in Fig. 1b, right panel).

Here, we describe a new method that builds on the single step principle to separate RNA, proteins and cross-linked RNA-protein complexes in biphasic extractions according to their physicochemical differences. Following the rationale that phenol and toluol (toluene) share a similar chemical structure but toluol is less water soluble due to the lack of the OH group (Fig. 1b), we modified the extraction chemistry using a mixture of phenol:toluol. This alters enrichment of the biomolecule classes in the extraction and furthermore enabled us to “shift" cross-linked RNPs into the aqueous or interphase, respectively (Fig. 1 b, left panel). Combining both separation strategies allowed us to sequentially deplete the sample of DNA and lipids, as well as non-cross-linked RNA and -proteins, highly enriching for cross-linked RNPs (clRNPs) that then can be directly analysed or further processed in more complex workflows.

## Results

### PTex enriches for cross-linked RNPs

The poly-A RNA interactome of human HEK293 cells has been mapped in great depth, providing an ideal reference to establish PTex-based (Phenol Toluol extraction) purification of cross-linked RNPs (2). After irradiation with UV light at 254 nm to induce covalent RNA-protein cross-links, cells were subjected to the PTex procedure, a series of three consecutive organic extractions:

- Step 1: Phenol:Toluol (PT; 50:50), pH 7.0
- Step 2: Phenol, pH 4.8, (chaotropic) detergents
- Step 3: Phenol, EtOH, water, pH 4.8

During the first extraction with Phenol/Toluol, RNA, proteins and cross-linked RNPs (clRNPs) are accumulating in the upper aqueous phase while DNA and membranes are predominantly found in the interphase (Fig. 1b left panel). The aqueous phase is subsequently extracted twice under chaotropic and acidic conditions using phenol (18). Now, free RNA accumulates in the upper aqueous phase, free proteins in the lower organic phase and clRNPs migrate to the interphase (4) (Fig. 1b right panel). Finally, the complexes in the interphase are precipitated using ethanol (19). To track the distribution of the diverse cellular molecules from total HEK293 cells during the purification procedure, we probed all phases from the intermediary steps by western blotting against HuR (ELAVL1), a well established 35 kDa RBP (20, 21) (Fig. 1c). UV-cross-linking produces an additional band at high molecular weight which indicates the RNA-cross-linked fraction of HuR (clHuR; Fig. S1). In the PTex fraction (interphase 3), clHuR was highly enriched whereas free HuR was strongly reduced. Furthermore, abundant cellular proteins unrelated to RNA-binding such as beta-actin are not detectable in the PTex fraction. To further demonstrate the efficient removal of non-cross-linked proteins, we spiked into the cell lysates a recombinant RNA-binding protein (the central domain of *Drosophila melanogaster* Sex-lethal, denoted Sxl-RBD4 (22)), after UV-cross-linking. PTex efficiently removes ~99% of the free spike-in RBP (as determined by densitometry compared to the input; Fig. 1c). Similarly, removal of “free" RNA was demonstrated in an *in vitro* assay in which 32P-5’ labeled RNA was subjected to PTex (Fig. 1d). We next tested for depletion of DNA by PCR targeting genomic DNA (exon 5 of the IL3 gene) or plasmid DNA (pUC19). DNA is removed during the first two PTex steps (Fig. 1e). We then tested for additional well-established RBPs (Fig. 1f), namely polypyrimidine tract binding protein 1 (PTBP1), fused in sarcoma (FUS), and the more recently identified RNA-binding enzymes glyceraldehyde-3-phosphate dehydrogenase (GAPDH) and enolase (Eno1)(3); all of which are enriched by PTex in a UV-irradiation-dependent manner whereas the highly abundant DNA-binding histone H3 is depleted. We additionally used PTBP1 and FUS to demonstrate that RNase digestion prior to PTex abolishes efficient enrichment of RBPs (Fig. 1g,S9), consistent with selectivity for RBPs complexed with RNA. In sum, PTex highly enriches for cross-linked RNPs while efficiently depleting non-cross-linked proteins or nucleic acids.

### PTex performance

We then set out to critically assess the performance of PTex to purify RBPs. However, while cross-linked HuR generates a signal at the height of the gel pocket (compare Fig. 1c and S1), we have no *a priori* knowledge about the amount of protein which is efficiently cross-linked by UV light to RNA. Thus, we performed RNA interactome capture(1) on HEK293 cells to obtain proteins which are 100% cross-linked to poly(A) RNA. We used this sample as input for PTex (Fig. 2a) and quantified purification of HuR by densitometry (Fig. 2b), resulting in a ~30% recovery of the cross-linked protein input. To analyse the minimal length of RNA required for PTex-mediated enrichment of RNPs, we again employed the recombinant and highly purified 20 kDa Sxl-RBD4 protein which associates with Uracil stretches of 7 nucleotides or longer (23). We produced *in vitro* transcribed RNAs with lengths varying between 13 and 191 nucleotides, all of which contained the same Sxl-binding motif at their 5’-end. After binding and UV cross-linking *in vitro*, samples were PTex-purified and analysed by western blotting (Fig. 2c). PTex efficiently recovered cross-linked Sxl-RBD4 complexes with RNA as short as 30 nt. Similar to HuR, we calculated the recovery of Sxl by densitometry (Fig. 2d) which is ~50% of the input cross-linked protein. Since purification efficiency differs for individual proteins (see Fig. 1c,1f, 2a-d), we quantified protein and RNA recovery by PTex using spectroscopy and measured absorption at 260 nm and 280 nm as proxy for RNA and protein, respectively (Fig. 2e). Consistent with the other results, overall PTex recovery is 27% (RNA) and 33% (proteins) from cross-linked HEK293 interactome capture samples (Table S4).

**Fig. 2.**
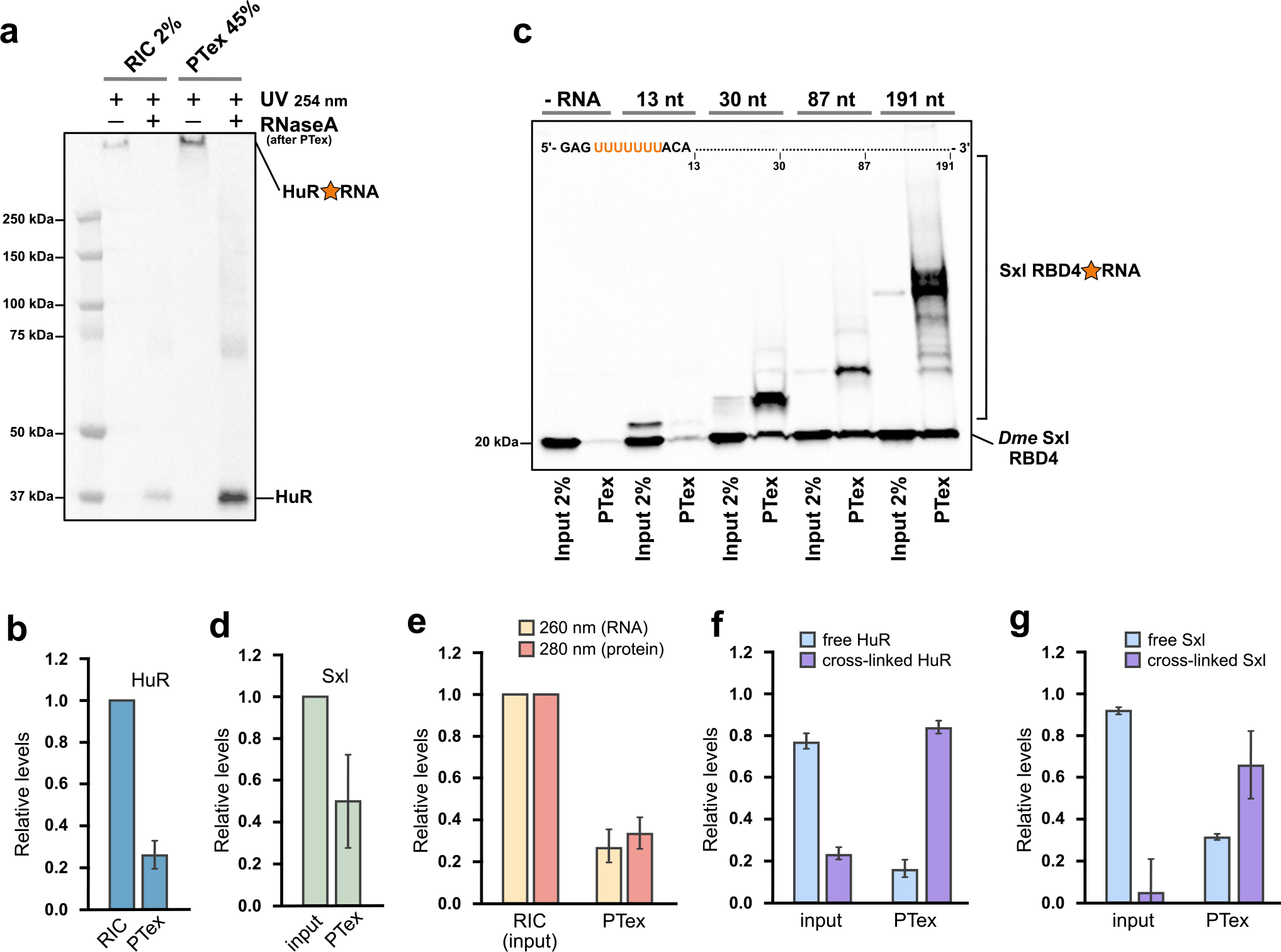
Performance of PTex. a) RNA interactome capture of HEK293 was used as 100% cross-linked input material and HuR recovery by PTex was assessed. b) Quantification of a (n=3). c) *Drosophila melanogaster* RBP Sxl RBD4 was bound to RNA of several length carrying the Sxl U7 recognition site and UV-cross-linked *in vitro*; minimal RNA length for efficient recovery by PTex is 30 nt. d) Quantification of c (n=3). e) PTex recovery of RNA (260 nm) and proteins (280 nm) determined by UV spectroscopy (n=3). f,g) Relative enrichment of cross-linked HuR (source Fig. 1c) and Sxl (source Fig. 2c, n=3) by PTex. All data are in Supplementary Table S4. For full blots see Supplementary Figures S10-S14.

Finally, we estimated the relative enrichment of free vs. cross-linked protein in input and PTex (Fig. 2f,g). For HuR (*in vivo*), PTex largely enriches for RNA-bound HuR with a “background” of 10-15% non-UV-cross-linked HuR. For Sxl, PTex-mediated enrichment is even larger if compared to the input; however, free Sxl is more prominently detected in the artificial *in vitro* conditions only but not when spiked-in to cell lysates (Fig. 1c) where the free protein is removed almost completely. Interestingly, we only recover non-cross-linked proteins in the PTex fraction when interrogating RNA-binding proteins but not in the case of other abundant cellular proteins such as histone H3 or beta-actin (see Fig. 1c,f,g). We attribute this to either stable RNA-protein complexes which resist complete denaturation and separation during the PTex procedure or an artifact of the gel system as RNA strand breaks after PTex purification could result in a western blot signal akin to the free/unbound RBP. In both cases, only bona-fide RBPs will be enriched.

### Purification of RBPs from animal tissue

To test if PTex can be directly applied to tissues, we UV cross-linked whole mouse brain samples and performed PTex to extract clRNPs directly (Fig. 3). Since brain tissue is particularly rich in lipids which accumulate in the interphase during step 1 of the PTex protocol, we increased the temperature during extractions to 65°C (Hot-PTex). The extracted clRNPs were analysed by western blotting. Cross-linked HuR is largely enriched over non cross-linked HuR after PTex (Fig. 3 lower panel). Weak detectable bands at 70 or ~110 kDa, respectively, are likely HuR dimers and trimers as observed before (24). This example demonstrates that PTex is not only suited for cell culture but can also extract RNPs directly from animal tissue samples; an advantage over purification protocols that depend on RNA labeling (such as PAR-CLIP(11), RICK(16) or CARIC(17)) for which efficient uptake and incorporation of nucleotide analogs can be challenging. Brain tissue can be efficiently UV cross-linked however as demonstrated by HITS-CLIP experiments (25).

**Fig. 3.**
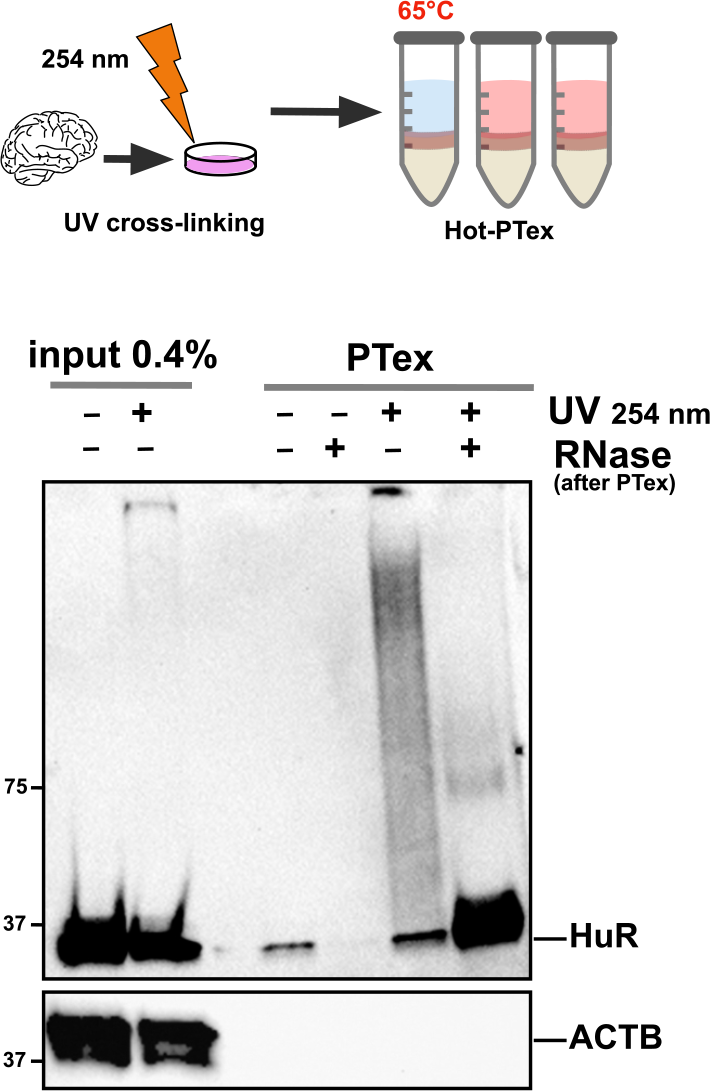
RNP purification from animal tissue. Mouse brain tissue was cryogrinded and UV-irradiated before (Hot)-PTex was performed. Western blot against HuR (ELAVL1) demonstrates recovery of cross-linked HuR from mouse tissue while beta-actin (ACTB) is efficiently depleted. For full blots see Supplementary Figure S15.

### A simplified CLIP protocol

HuR (ELAVL1) has been shown to interact with mRNA and pre-mRNA in several CLIP studies and has a well-documented binding motif (5’-UUUUUU-3’) (26). After *in vivo* labeling of cellular RNA using 4-thiouridine (4SU) and UV irradiation at 365 nm, we performed i) classical PAR-CLIP analysis (PAR-CLIP-classic)(11, 27) of HuR, ii) a PAR-CLIP variant using on bead ligation of adapters (PAR-CLIP-on-beads)(28, 29), and iii) a version in which we use phenol extraction (termed pCLIP) for removal of unbound RNA instead of PAGE/membrane excision (Fig. 4a). We found that pCLIP libraries contained a larger fraction of longer reads than the PAR-CLIP classic/PAR-CLIP-on-beads libraries (Fig. 4b). All three approaches identify the canonical 5’-UUUUUU -3’ motif and similar profiles of HuR-bound RNA clusters map to intronic and 3’UTR regions (26) in all three variants (Fig. 4c-e). The clusters could also be mapped to the same 3’UTR loci when comparing HuR binding sites in tubulin and splicing factor Srsf6 mRNA (Fig. 4f,g). Although we only performed a low-read-coverage experiment as proof-of-principle, our results demonstrate that phenolic extractions of RNPs such as PTex can be integrated into more complex workflows such as (PAR-)CLIP and have the potential to simplify CLIP-type approaches by enriching for clRNPs or remove unbound RNA transcripts.

**Fig. 4.**
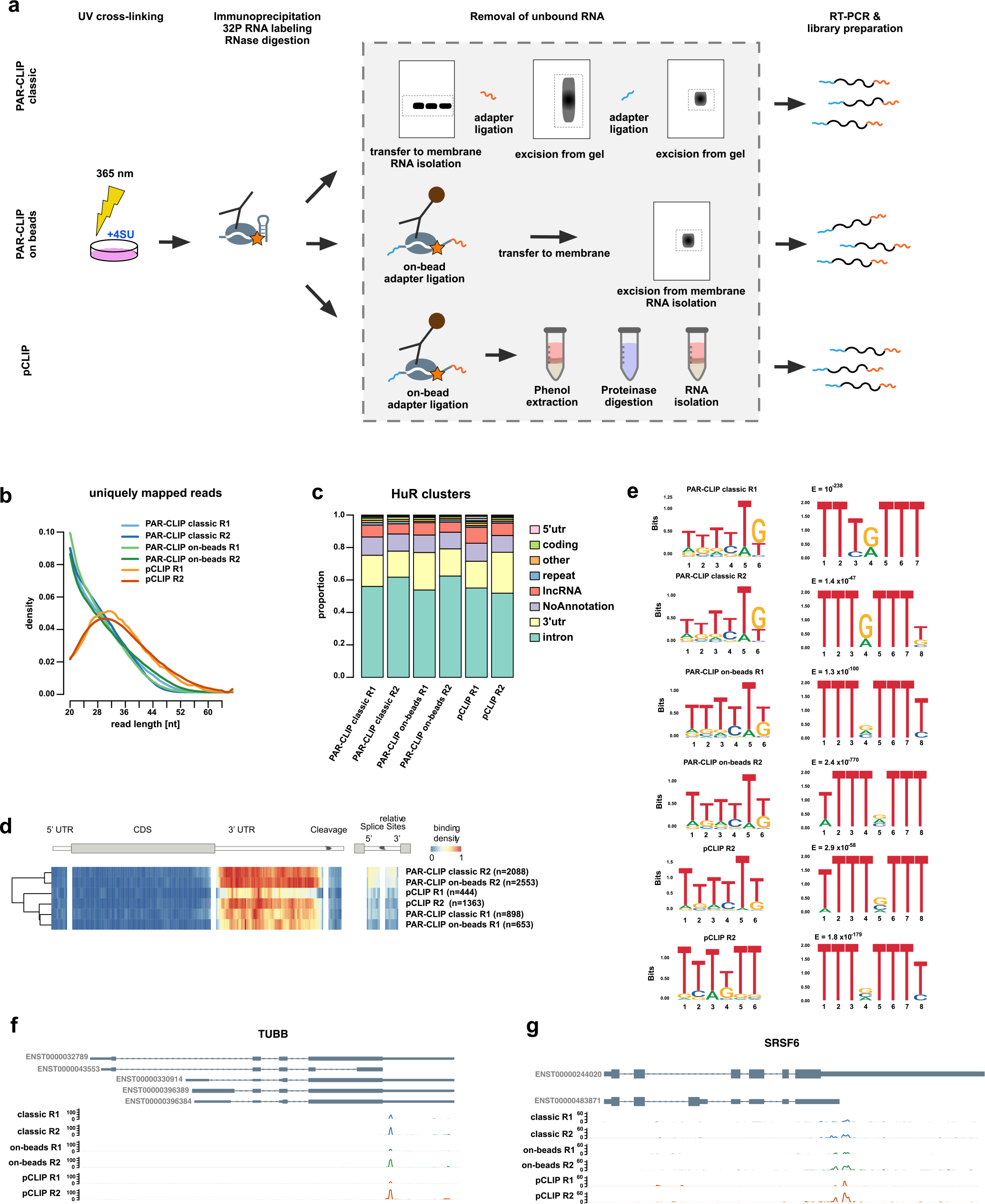
pCLIP: a fast PAR-CLIP variant employing phenolic extraction. a) Schematic comparison of PAR-CLIP variants. b) Read length distribution of uniquely mapping reads utilized for determine binding sites (cluster) of HuR (ELAVL1). PAR-CLIP samples were processed using PARpipe (see methods). c) Relative proportion of PARalyzer-derived cluster annotation. d) Heatmap of relative positional binding preference for intron-containing mRNA transcripts for each of the six HuR PAR-CLIP samples. Sample-specific binding preferences were averaged across selected transcripts (see methods). The relative spatial proportion of 5’UTR, coding regions and 3’UTR were averaged across all selected transcript isoforms. For TES (regions beyond transcription end site), 5’ splice site, and 3’ splice site, we chose fixed windows (250 nt for TES and 500 nt for splice sites). For each RBP, meta-coverage was scaled between 5’UTR to TES. The 5’ and 3’ intronic splice site coverage was scaled separately from other regions but relative to each other. e) We applied de novo motif discovery for PARalyzer derived clusters using ZAGROS (left) and DREME (right). For Zagros(30), we found a T-rich motif scoring the highest in all cases. As ZAGROS does not return E-values we analyzed the cluster sequences using DREME. For all but classic PARCLIP R2 we found a T-rich motif scoring the highest. For classic PARCLIP R2 however, the T-rich motif scored second with a similar E-value to a less frequent primary motif (Supplementary Fig. S16). f) and g) Genome browser shots of TUBB and SRSF6 example genes showing reproducible 3’UTR binding sites. Track y-axes represent uniquely mapping read count.

### A global snapshot of human RNA-protein complexes

Despite the recent advances in mapping RBPs in many species, two general issues have not been addressed to date: i) the fact that RNA interactome capture (1–3) targets only polyadenylated RNAs suggests that many RBPs that bind non-adenylate RNAs are missed by this experimental approach (16); and ii) although UV cross-linking has been widely used to research RNA-protein interactions, no systematic study has been performed to determine the optimal irradiation conditions for efficient cross-linking of individual RNPs.

Using PTex as an unbiased approach, we set out to explore RNA-protein interactions cell-wide in HEK293 cells. To test the effect of different energy UV-irradiation, we employed besides the most commonly used 0.15 J/cm^2^ irradiation at 254 nm wavelength also irradiation at two additional energy levels: 0.015 and 1.5 J/cm^2^; spanning two order of magnitude of UV irradiation dosage. This setup was then used to comprehensively map RNA-protein interactions beyond the established poly-A RNA-bound proteomes (2, 8). We independently irradiated HEK293 cells at all 3 energies and performed PTex purification of cross-linked and non-cross-linked RNPs from whole cells (Fig. 5a). Transcriptomics by RNA-Seq (Fig. 5b,c), and proteomics by mass spectrometry and label free quantification (LFQ) (Fig. 5d-g) were performed using total RNA and protein preparations as input controls.

**Fig. 5.**
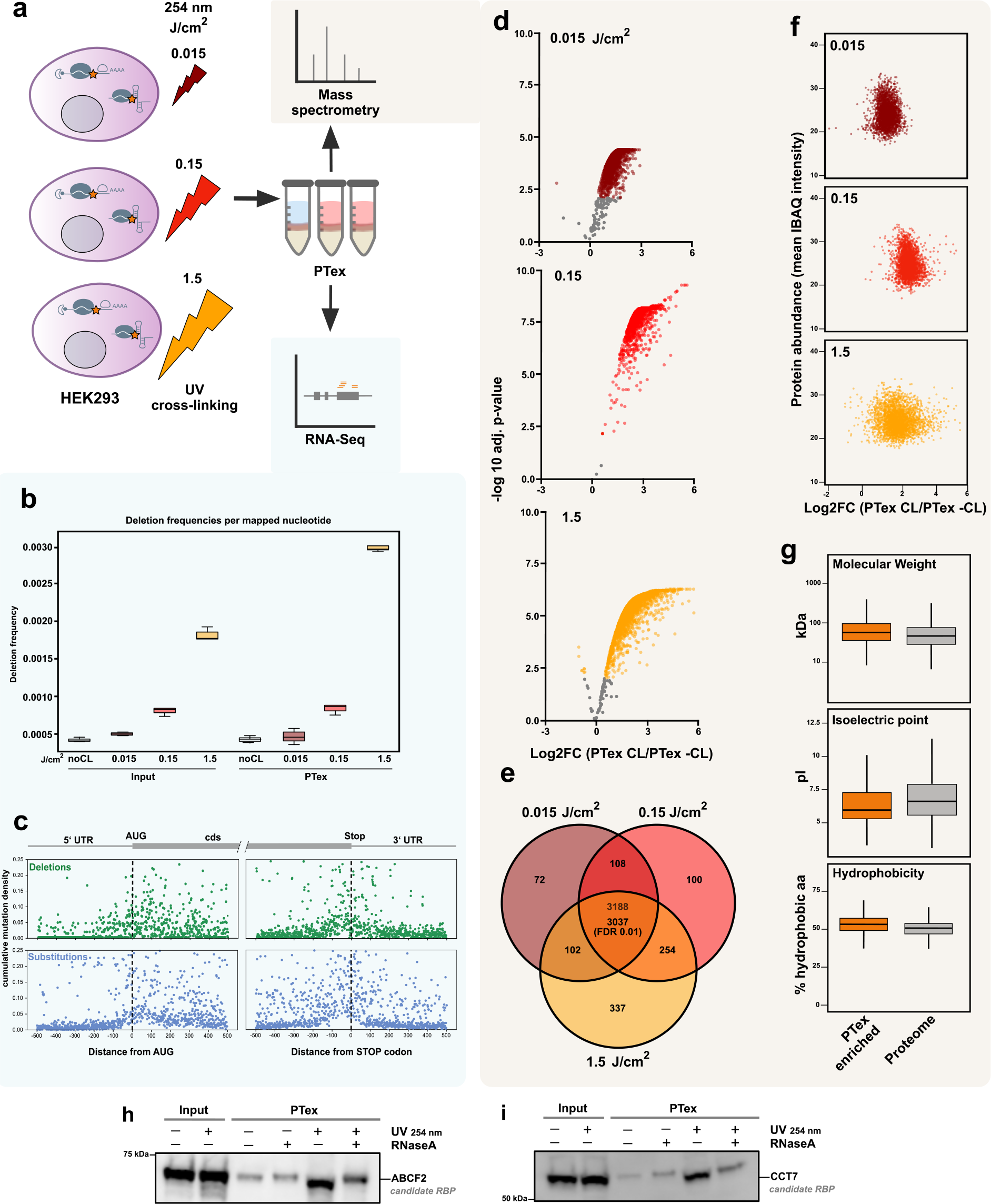
A global snapshot of RNPs in HEK293 cells (RNA) a) Schematic of the experimental setup: HEK293 cells were UV-cross-linked using 0 (noCL),0.015 (dark red), 0.15 (red) and 1.5 (dark yellow) J/cm^2^ 254 nm light in triplicates. Total RNA from input (whole cell lysate) and PTex-purified samples were analysed by RNA-Seq. b) Deletions in RNA from input and PTex samples; frequency of mutations in transcripts correlate with higher UV doses. c) Mutations (deletions = green, substitutions = blue) enriched in UV-irradiated samples were plotted to their position relative to AUG and Stop codon in coding sequences and serve as indicator for protein-binding sites. Note that we cannot delineate which protein bound to which position. Plots for PTex are shown; for input see Supplementary Fig. S17,18. d-g) Input (whole cell lysate) and PTex-purified sample were analysed by label-free mass spectrometry. d) Volcano plots of proteins enriched by PTex (FDR 0.01) under the three cross-linking conditions. e) Overlap of PTex-enriched proteins (enriched in all 9 replicates, FDR 0.01) is 3037 (these PTex proteins are from here on colored in orange). f) Protein abundance (IBAQ intensities of input samples) does not correlate with PTex enrichment (log2-fold change of intensities [CL/-CL]) g) PTex does not select for a subset of proteins based on general features such as molecular weight, pI or hydrophobicity. f) Enrichment of proteins by PTex drops at 1.5 J/cm^2^ 254 nm light compared to lower doses. h,i) PTex of individual predicted RNA-associated proteins. ATP-binding cassette sub-family F member 2 (ABCF2) and T-complex protein 1 subunit eta (CCCT7) have not been reported to bind RNA. Both are enriched after PTex in a UV-irradiation-dependent fashion, indicating that they indeed associate with RNA *in vivo*. For full blots see Supplementary Figure S19.

### PTex-purified RNPs:RNA

We first analysed PTex-purified transcripts. Unlike proteins which can be grouped into RNA-interactors and non-interactors, all cellular RNA can be expected to be associated to proteins (31–33). In line with this, we find a similar distribution of RNA classes when comparing RNA from inputs and PTex by RNASeq with the vast majority of transcripts being ribosomal RNA (Fig. S21). Protein-cross-linked RNA is known to enrich for mutations during reverse transcription in the RNASeq workflow (Fig. 5b,S17) (10, 11). Such mutations can then be used as “beacons” to map protein binding sites in transcripts (2, 11, 34). We used pyCRAC (35) to map deletion and substitution mutations enriched in UV-treated PTex samples to 100 nt windows around the 5’ AUG start codon and 3’ Stop codon of mRNA reads (Fig. 5c,S18). We find that most mutations (indicating protein binding) are within the first 100 nt after the AUG and the last 100 nt before the Stop codon. This was observed before in a global protein occupancy profiling study (36) and can potentially be attributed to cross-linking of ribosomal proteins or translation initiation/termination factors to mRNA, as ribosomal profiling experiments show increased ribsomal footprint densities at these regions, indicating longer dwell times and a higher potential for cross-linking at these sites (37).

### PTex-purified RNPs: proteins

Proteins which were not identified by MS in all 3 replicates after PTex were removed. For the remaining proteins, ratios of cross-linked over noncross-linked (CL/-CL) LFQ intensities (from the PTex experiments) were calculated (Fig. 5d). P-values from a moderated t-test were then used for multiple testing (BenjaminiHochberg). Using these stringent tests, we identify 3188 shared among the three conditions; out of these, 3037 proteins are significantly enriched in a UV-irradiation-dependent fashion in all samples using a cut-off of FDR 0.01 (Fig. 5e, Supplementary Table S1). Analysis of general protein features (molecular mass, pI, cellular abundance, hydrophobicity, Fig. 5f,g) demonstrates that the PTex procedure does not enrich for a particular subgroup of cellular proteins based on chemical properties or expression level. We picked two of the 3037 proteins which have not been reported to bind RNA: ATP-binding cassette sub-family F member 2 (ABCF2), a member of the AAA+ ATPase family (see below) and T-complex protein 1 subunit eta (CCCT7) as part of the chaperonin CCT/TRiC which is involved in telomere maintenance (38). Both are enriched after UV-irradiation *in vivo* and PTex purification (Fig. 5h,i), indicating RNA-association.

In principle, extended UV exposure should increase the chance for cross-linking events and thus for subsequent protein recovery by PTex (3). We therefore expected a gradual increase in enrichment of RNA-interacting proteins after PTex with higher UV dose. However, we only find increased recovery for most proteins when comparing low and medium energies. Surprisingly, irradiation with 1.5 J/cm^2^ on the other hand leads to a significant decrease in recovery (Supplentary Fig. S23). We observed RNA degradation when analysing total RNA from HEK293 cells irradiated with 1.5 J/cm^2^ 254 nm UV light (Fig. S20,S21; (3)). If this degradation was due to damage of nucleic acids induced by high energy UV light or a result of secondary processes during the extended time of treatment is unclear; in any case extensive shortening of RNAs will cause a loss of RNPs purified by PTex.

### HEK293 RNA-interacting proteins

So far, 700 - 2000 well established and recently identified eukaryotic RBPs have been described (reviewed in (39) and (8)) and PTex-purified protein patterns differ from the whole proteome in silvers stains (Supplementary Fig. S22). However, to find more than 3000 proteins to be enriched as RNA-associated by PTex is unexpected. Considering that deep proteome studies detect around 10,500 proteins in HEK293 cells (40), we find nearly a third of the expressed cellular proteome to be associated with RNA.

To test sensititvity and specificity of our approach, we first performed global GO enrichment analysis showing that terms from all aspects of RNA biology are the most enriched among PTex-purified proteins (Fig. 6a). At the same time, protein classes with no general role in RNA biology such as transporters and (trans-)membrane proteins were depleted by PTex (Supplementary Table S2). Known RNA-binding domains (RBDs) such as the RNA recognition motif (RRM), helicase folds (DEXDc, HELICc) or K homology (KH) domain were significantly enriched among PTex-purified proteins (Fig. 6b; Table S2). 89 PTex-enriched proteins contain a WD40 fold; a domain found to directly bind snRNA in Gemin 5 (41), taking part in rRNA biogenesis (Erb1 (42)) and found in RBPs (1). Other enriched domains are: AAA (AT-Pase, see below) fold, tetratrico peptide repeat region (TPR) as found in the yeast Clf1p splicing factor (43), Ski complex (44) and the translation terminator Nro1 (45), and the CH domain which is found among actin-binding proteins (3).

**Fig. 6.**
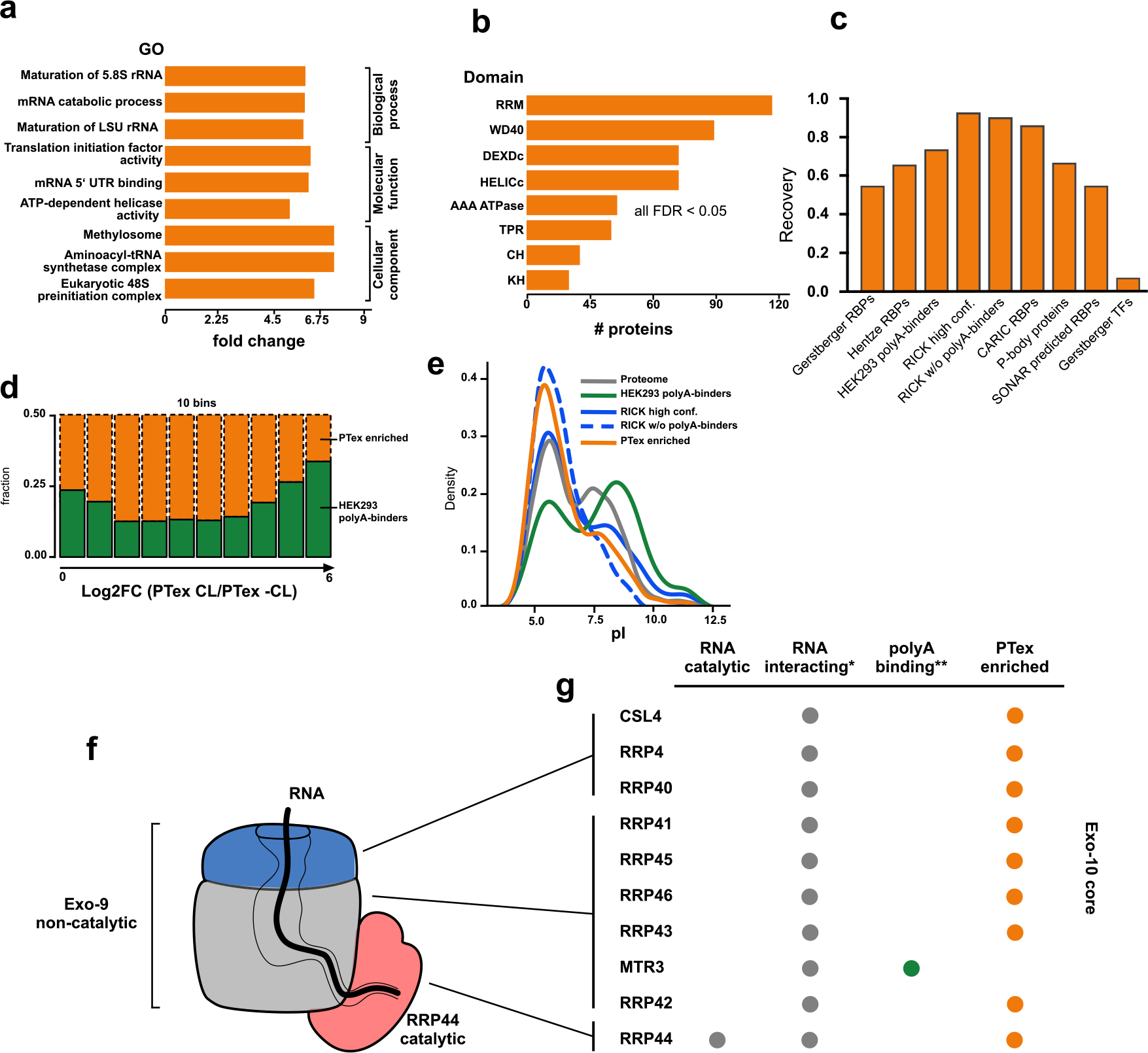
Features of RNA-interacting proteins found by PTex. a) Top 3 enriched GO terms (CC, MF, BP) and b) enriched protein domains in PTex-purified proteins from HEK293 cells. c) PTex-purified proteins overlap with well-described RBPs but not transcription factors. Recovery of Gersberger RBPs and transcripiton factors (TFs) reviewed in (39), a recent review on RBPs by (8), HEK293 poly-A binders (2), RBPs found by RNA interactome using click chemistry (RICK(16); CARIC(17)), P-body components (56) and a recent prediction of candidate RBPs (SONAR, (57)). d) Distribution of previously identified HEK293 mRNA-binding proteins (green; (2)) in PTex; each bin represents 10% of the 3037 PTex proteins from lowest to highest enrichment. e) mRNA-binding proteins display a bimodal pI distribution pattern with peaks at pH 5.5 and 9.5 (1, 2). RNA-interactors in general peak at pI 5-6 as found by PTex and RICK (16). **Proteins of the RNA exosome are prototype PTex proteins.**f) The RNA exosome core consists of ten subunits: nine non-catalytically active proteins (Exo-9) forming a barrel-like structure and an additional RNase (Rrp44; Exo-10); modified from (50). g) *All exosome subunits are labeled “RNA-binding” (Uniprot.org); **green = identified via poly-A selection in (2); orange = enriched in PTex.

To test that we are enriching for RNA-binders specifically, we calculated the probability of recovering known RBPs interacting with different RNA classes using the hypergeometric test: we enrich for ribosomal proteins (42/47 large subunit and 30/33 small subunit (46); p-value: 1.36 × 10^−30^), NSUN2 and tRNA synthetases (19/20 cytosolic; p-value: 7.4 × 10^−10^) indicating that our approach indeed captured cellular RNPs in a poly-A-independent fashion. We recover 70% of poly-A RNA-binding proteins found in HEK293 cells by (2) (Fig. 6c) although different UV irradiation strategies were used (254 vs. 365 nm cross-linking; p-value: 1.26 × 10^−139^). Importantly, the largest overlap was with RBPs recently found in HeLa cells using the also unbiased RICK technique (16) (94% for high confidence RBPs and 86% of non-poly-A RBPs, respectively) and the CARIC approach (84% overlap)(17) (Fig. 6c). Recent studies show that the boundaries between RNA- and DNA-binding are rather blurry and nuclear DNA-binders were found to interact with RNA (47). We identified proteins involved in replication and response to DNA damage (Table S2) such as DDX54 (48) but in general, DNA-binders such as transcription factors were underrepresented, demonstrating that PTex does not select for DNA-specific binding proteins in particular (Fig. 6c). To rule out that previously described RBPs are more efficiently recovered in PTex than the newly identified RNA-associated proteins (which could indicate carry-over of proteins unrelated to RNA interactions), we compared the distribution of the HEK293 mRNA-binding proteins (2) in the PTex enrichment. The established RBPs are similarly enriched along the dynamic range (from no enrichment to log2FC PTex [CL/−CL] of 6; Fig. 6d) of PTex and hence display no difference to the novel RNA-interactors.

The presented results demonstrate that PTex is specific for RNPs. But why weren’t the same proteins discovered to associate with RNA before? The majority of recently discovered RBPs are interacting with mRNA (8); a RNA class which is highly heterogeneous in its sequence but represents only ~5% of the cellular RNA pool. The differences in between interactome capture (poly-A RNPs) and PTex (RNPs in general) is best demonstrated in the case of the eukaryotic RNA exosome (49, 50). The core exosome complex consists of ten protein subunits (Exo-10) from which only one protein (Rrp44) is catalytically acting on RNA as exo- and endoribonuclease (Fig. 6f). The remaining nine proteins (Exo-9) are forming a barrel-like complex in which RNA can be channelled through before it is degraded by Rrp44, but Exo-9 proteins do not degrade or modify the RNA itself (50, 51). Still, all ten subunits are positioned to directly interact with RNA and multiple interactions with the individual subunits have been demonstrated in high resolution structure studies (52–55). Hence, all ten subunits are amenable to UV-cross-linking and, as a result, 9 out of the 10 subunits were identified by PTex (Fig. 6g).

Previously it was demonstrated that interactome capture enriches for mRNA-binding proteins with a high isoelectric point (pI; Fig. 6e) (1, 3). However, sequence- and oligo-dTindependent approaches such as RICK (16) or PTex identify more proteins with a pI <6. Proteins with a low pI are overall negatively charged at cellular pH and thus unlikely to interact with RNA in an unspecific manner due to electrostatic repulsion of likewise negatively charged RNA. Indeed, 7 of the Exo-10 protein subunits have the mRBP-untypical isoelectric point below pH 6 (Supplementary Fig. S24). Inside the central exosome channel, RNA of 30-33 nt or 9-10 nt length has been found in *in vitro* and in CRAC analyses (58); the former being of sufficient length for efficient recovery by PTex (Fig. 2c). In sum, PTex enriches for the (near) complete exosome core complex while interactome capture from the same cell line only found a single subunit (2) (Fig. 6g).

Yet we also enrich for proteins which have no established role in RNA biology such as subunits of the human proteasome (Table S2). However, from the 28 proteins of the 20S core complex, Psma5 and Psma6 were reported to display RNase activity in purified complexes (59). Importantly, proteasome-associated RNAs were shown to lack poly-A stretches (60, 61) and ATPases of the AAA family in the 19S proteasome regulatory particle were found to be recruited to RNA polymerase I (rRNA) transcription sites (62). Hence, none of the proteasome-related RNA activities are approachable via poly-A RNA-mediated purification. The proteasome is a multi-protein complex with structural similarities to the exosome (50) and RNAs interacting with Psma5/6 are likely to be UV-cross-linked to other subunits as well.

### PTex allows a first snapshot of RNA-associated proteins in bacteria

Proka ryotic mRNAs lack poly-A tails and are thus not approachable by oligo-dT-based methods. We used the pathogen *Salmonella* Typhimurium harbouring a chromosomally FLAG-tagged Hfq protein (63) to test PTex in bacteria. Hfq is an abundant RNA-binder facilitating mRNA:ncRNA interactions in Gram-negatives (63, 64). As for animal tissue culture (see above), we used the slightly modified PTex protocol (Hot-PTex) in which RNP extraction from *Salmonella* grown to OD_600_ 3.0 was performed at 65°C thereby supporting cell lysis (Fig. 6a). Hfq is a 17 kDa protein and forms a homo-hexamer in bacterial cells; the complex has been shown to resist normal Laemmli PAGE conditions (63) and is also visible in western blots (Fig. 6b). After UV irradiation, PTex purification and RNase treatment, we observe a shifted Hfq monomer band which we attribute to residual cross-linked RNA fragments resulting in a slightly higher molecular mass. The physiologically relevant Hfq hexamer is also strongly enriched compared to non-UV samples, indicating that also the complex is still bound to remaining RNA fragments.

We next used the Hot-PTex fraction of *Salmonella* cells to map RNA-associated proteins by mass spectrometry (Fig. 6c). Comparing recovered protein intensities from UV irradiated versus control cells (biological duplicates), we find 172 proteins (Supplementary Table S3), among them 33 ribosomal proteins, components of the RNA polymerase complex (subunit *α*, *σ* factor RpoD, DksA) and 4 out of the 5 established mRNA-binding proteins of *Salmonella* (Hfq, ProQ, CspC/CspE) (63, 65, 66). 113 of the enriched proteins are so-far unknown to interact with RNA. To validate our findings, we picked YihI, a putative GTPase-activating protein which was speculated to play a role in ribosome biogenesis (67), the alkyl hydroperoxide reductase c22 protein (AhpC)(68) and the cell invasion protein SipA (69, 70). Using mutant strains carrying FLAG-tags fused to the C-terminus of PTex candidate RNA-interactors, we performed UV-cross-linking, immunoprecipitation and radioactive labeling of co-purified RNA (known as “PNK assay” as described in (71)), validating that YihI, SipA and AhpC are indeed associated to RNA *in vivo* (Fig. 6d,S25,S26). Additionally, we recently validated PTex-enriched ClpX and DnaJ as well (72). We furthermore find proteins with known RNA-binding domains (RBDs) such as the nucleic acid binding OB-fold (present in RpsA, RpsL, RplB, CspC, CspE, Pnp, RNaseE, Ssb, NusA) and domains which were also detected in RBPs when screening eukaryotic cells: the afforementioned AAA ATPase fold in the ATP-dependent protease ATPase subunits HslU, ClpB and ClpX, or thioredoxin domains as in AhpC, Thiol:disulfide interchange protein (DsbA) and Bacterioferritin comigratory protein (Bcp) (1, 3). As in many other species, we find glycolytic enzymes to associate to RNA (Pgk, Pgi) (3, 73). Using GO terms for functional annotation of the RNA-associated proteins, the most significant terms are “translation” and other terms connected to the ribosome as expected (Fig. 6e). To the best of our knowledge, the periplasmic space is generally considered to be devoid of RNA. We still recover RNA-associated proteins which localise to the outer membrane (Table S3). As in the case of HEK293, we cannot distinguish here between RBPs that actively act on RNA and proteins which are associated to RNA for e.g. structural reasons. However, recent studies in several Gram-negative bacteria demonstrate that secreted outer membrane vesicles (OMVs) contain RNA which indicates that bacterial transcripts must be sorted to the outer membrane via a yet to be determined pathway (reviewed in (74)). It is tempting to speculate if the here identified proteins are involved in such a process or are mere bystander proteins.

Overall, we noticed that enrichments even from known RBPs were lower in *Salmonella* compared to HEK293 cells and we anticipate that additional modifications in UV-cross-linking and/or cell lysis could improve sensitivity when applying (Hot-)PTex in bacteria. PTex is the first approach that can purify bacterial RNPs in an unbiased fashion without the necessity of immunoprecipitation or introduction of modifications (tag, overexpression, etc.), rendering it a novel tool for cell-wide RBP identification and studying bacterial RNA-protein interactions.

## Discussion

UV cross-linking of RNPs appears rather inefficient. Even after high UV doses only ~1-10% of any given RBP can be covalently coupled to ribonucleic acids in human cell culture (1, 4) and yeast (3, 10, 73, 75). Importantly: this reflects both cross-linking efficiency and fraction of protein bound to RNA; in other words how much of the protein is in a steady state associated with the RNA? What is of experimental interest is therefore only a minor fraction of the RBP (the one cross-linked to RNA), while the vast excess of protein stays in a non-cross-linked state.

With PTex, we are exploiting the physicochemical differences between cross-linked hybrid RNA-protein molecules on the one hand and the non-cross-linked proteins and RNA of a cell on the other for selective purification of complexes only. The method was designed to select against secondary RNA-binders by using denaturing and chaotropic conditions in which RNA-protein interactions which were not covalently cross-linked are not preserved (18), and by selecting proteins which were enriched in a UV-dependent fashion. PTex is a fast and simple modular protocol which can be performed in about 3 hrs. Our approach is independent of the UV wave length applied for irradiation (254/365 nm) and the type of biomaterial used (human/bacterial cell culture, animal tissue), and does not rely on presence of a particular RNA sequence such as poly-A tails. 30-50% of cross-linked proteins could be recovered by PTex compared to the starting material (Fig. 2b-e). At the same time, PTex drastically improves the relative enrichment of cross-linked over free protein (Fig. 2f,g).

Next to PTex, RBR-ID(76), RICK(16) and CARIC(17) are methods which allow for unbiased purification of RNPs. However, there are also differences: All, RBR-ID, RICK and CARIC rely on RNA labeling using either 4SU or 5-ethynyl uridine (EU) and subsequent UV irradiation. Using pulse/chase experiments, these methods thus allow to determine newly-transcribed RNA species. For PTex, UV irradiation at 254 nm wavelength is sufficient (no labeling required) and we anticipate that this is an advantage when analysing biological material in which uptake of nucleotide analogs is either insufficient or too cost-intensive. It did not escape our attention however that PTex requires fewer input material compared to the other methods (Table 1). Since all of these methods rely on UV irradiation however, biases introduced by different cross-linking efficiencies for individual proteins remain a general issue.

**Table 1.** Starting material of RNP purification methods.

| Method | Starting material (cells) | Study |
| --- | --- | --- |
| **PAR-CLIP** | $2 - 9 \times 10^8$ | (11) |
| **RIC** | $2.85 \times 10^8$ | (1) |
| **RBR-ID** | N/A | (76) |
| **RICK** | $2 \times 10^7$ | (16) |
| **CARIC** | $3.6 \times 10^8$ | (17) |
| **PTex** | $5 - 8 \times 10^6$ | this study |

Our work provides a cell-wide analysis of the effects of different UV irradiation dosage on RNA-protein cross-linking (Table S1, Figure S23). We hope that this resource will aid researches to establish suitable conditions for cross-linking of individual RNPs. The decrease in recovery of proteins after using 1.5 J/cm^2^ 254 nm light (Fig. S23) demonstrates that extensive irradiation/cross-linking can have adverse effects on protein recovery. Next to the observed RNA degradation, cross-linked peptides released by tryptic digestion are notoriously difficult to identify in MS experiments and hence increasing the amount of protein cross-linking might negatively impact protein identification (76–79). This is not the case for PTex-purified and input control transcripts in which deletion mutations accumulate at high UV settings which can then serve as marker for protein interaction sites (Fig. 5b,c) (10). Our results demonstrate that when investigating RNPs on a global scale, the cross-linking strategy should be adapted to the biological question: not all proteins are interacting with RNA and increasing the UV dose can be disadvantageous for RBP recovery since severely degraded RNA will cause less-efficient purification by PTex and cross-links will impair identification by mass spectrometry. Conversely, almost all RNA can be expected to be bound by a set of proteins under physiological conditions (32, 33) which explains why we also observe an increase in mutations in input RNA (Fig. 4b,S17,S18). In contrast to protein recovery, (partial) *in vivo* RNA degradation will not impair recovery of cross-linked transcripts since RNase treatment/RNA fragmentation is part of CLIP and RNASeq workflows already and cross-linked RNA will not be lost during cDNA preparation or sequencing (9–15).

Our findings indicate that up to a third of a cell’s proteins can associate with RNA *in vivo* which raises the question of the underlying biological function of these interactions. In this respect, it is intriguing to see that the Exo-9 proteins are interacting with and can be cross-linked to RNA although none of these subunits display RNase activity themselves. By increasing the detection efficiency for UV-cross-linked complexes e.g. by recovery of proteins interacting with RNA as short as 30 nt, we now have to separate classical RBP functionalities such as RNA degradation, transport or modification from RNA-interactors which are in physical contact with RNA due to structural organisation as in the case for ribosomal or exosome proteins. With PTex, we have developed a tool for fast recovery of RNPs in general which will allow us interrogate the functionality of individual proteins in RNA biology. So far, eukaryotic proteomes have been extensively scrutinised for RNA-binding proteins in the recent years. The two other kingdoms of life - archea and prokaryotes - could not be investigated for technical reasons. We here provide a first RNA-bound proteome from *Salmonella* Typhimurium, demonstrating that PTex will now allow to expand global RNP analysis to species in all three branches of the tree of life.

## Methods

### Human cell cell culture and *in vivo* cross-linking

Human embryonic kidney cells (HEK293) were grown on 148 cm^2^ dishes using DMEM high glucose (Dulbecco’s Modified Eagle, glucose 4.5g/L, Gibco, 41966-029) supplemented with 10% bovine serum (Gibco, 10270-106), penicillin/streptomycin (100 U/mL 0,1 mg/mL; Gibco, 15140-122) at 37°C with 5% CO2. After reaching 80% confluence, cells in monolayer were washed once with cold phosphate buffer saline (DPBS; Gibco, 10010-015) and placed on ice. Then, DPBS was removed completely and cells were irradiated with 0.015 − 1.5 J/cm^2^ UV light (λ= 254 nm) in a CL-1000 ultraviolet cross-linker device (Ultra-Violet Products Ltd), collected in 15 mL tubes, pelleted by centrifugation (1000 × g, 3 min, 4 °C), aliquoted in 2 mL tubes and stored at −20/-80°C (+CL). Non-irradiated cells were used as non-cross-link control (−CL). Additionally, the potential UV damage on the RNA after exposure to the different radiation energies was assessed: RNA isolated from HEK293 cells before and after exposure UV light by phenol extraction (18) were analysed with the RNA 6000 Chip Kit in a Bioanalyzer 2100 (Agilent, 5067-1513).

### Bacterial cell culture and *in vivo* cross-linking

*Salmonella* Typhimurium Hfq::x3FLAG (63) was grown on LB medium to stationary phase (ODM_600_ = 3). Aliquots of 20 ml were pelleted (20,000 × g, 8 min, 37°C) and resuspended in 1/10 of water for UV irradiation. Cells were cross-linked on ice with 5 J/cm^2^ UV light (λ= 254 nm) in a CL-1000 ultraviolet cross-linker device (Ultra-Violet Products Ltd), snapfrozen and stored at −80°C. Bacterial suspensions equivalent to 2.5 ml of initial culture were used as input in figure 7b. *Salmonella* enterica subsp. enterica Serovar Typhimurium strain SL1344 used for the mapping of RNA-protein interactions was grown in LB medium to OD_600_ 2.0. Half of the cultures were cross-linked in a Vari-X-linker (UVO3, http://www.vari-x-link.com), using UV light (λ= 254 nm) lamps for 90 seconds. Fractions of 10 ml from each, cross-linked and non-cross-linked cultures, were harvested by filtration as described in (80).

**Fig. 7.**
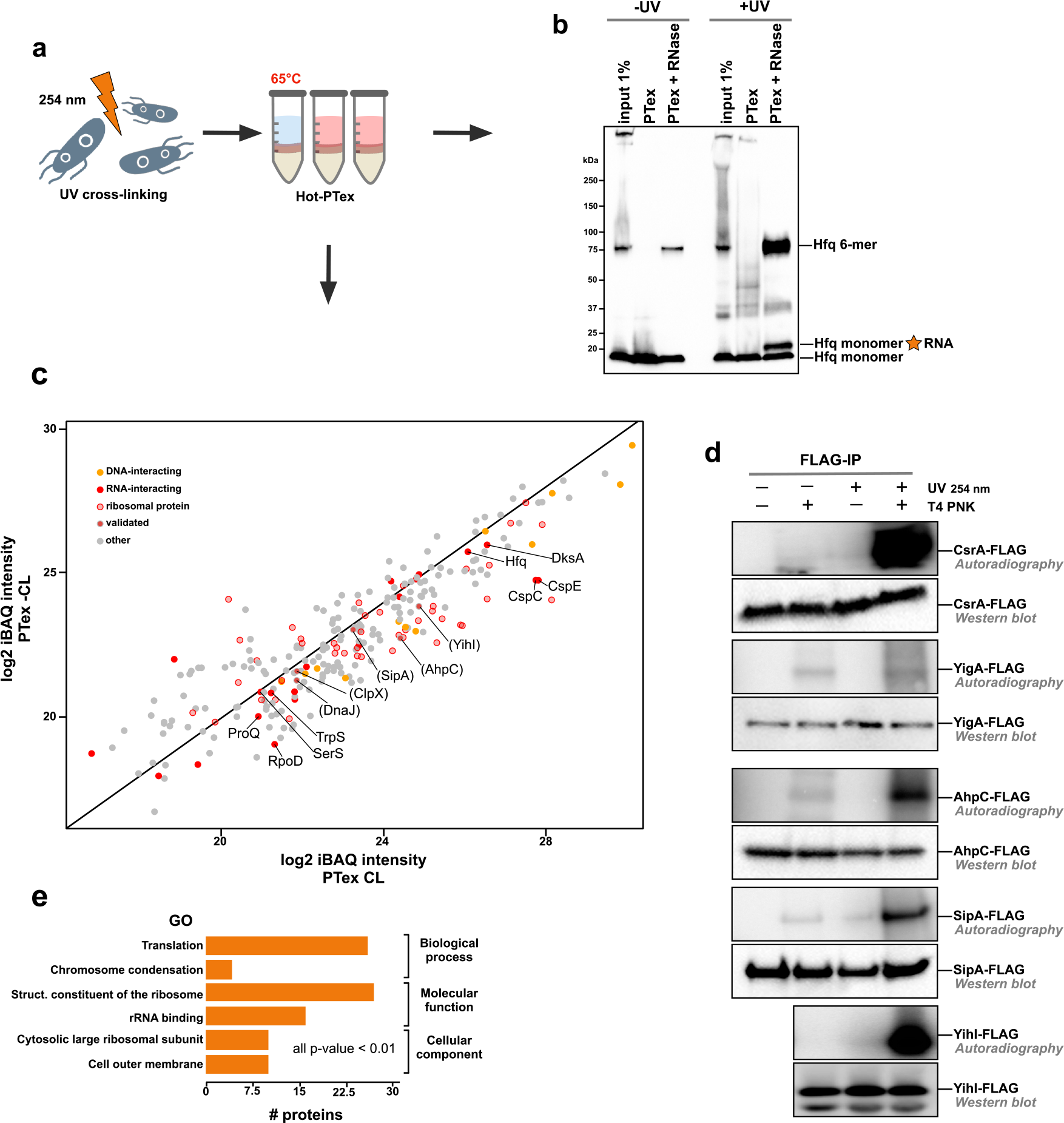
PTex recovers bacterial RNPs. a) *Salmonella* Typhimurium SL1344 Hfq-FLAG was UV-cross-linked and HOT-PTex was performed to purify bacterial RNPs. b) Western blot using an anti-FLAG antibody demonstrates recovery of Hfq monomers linked to RNA. Note that the physiologically active Hfq hexamer partially withstands SDS-PAGE conditions (63) and that this complex is also enriched after PTex. c) RNPs in *Salmonella* were purified by PTex globally. 172 Proteins enriched after UV-crosslinking (PTex CL) contain ribosomal proteins (transparent red), known RBPs (red) and DNA-binders (orange). Individual enriched proteins not known to associate with RNA before were used for validation (in parentheses). d) Validation of PTex-enriched RNA-interactors: *Salmonella* strains expressing FLAG-tagged proteins were immunoprecipitated +/− UV irradiation. RNA-association is confirmed by radioactive labeling of RNA 5’ ends by polynucleotide kinase (T4 PNK) using autoradiography; a signal is exclusively detectable after UV-crosslinking and radiolabeling of precipitated RNA. CsrA-FLAG (pos. ctr.), YigA-FLAG (neg. ctr.), AhpC-FLAG, SipA-FLAG and YihI-FLAG are bound to RNA *in vivo*. e) GO terms significantly enriched among the RNA-associated proteins. For full gels/blots see Supplementary Figures S25,S26

### Construction of bacterial strains

Yihi::x3FLAG::KmR was constructed following the procedure based on the Lambda Red system developed by (81). The system is based on two plasmids: pKD46, a temperature-sensitive plasmid that carries gamma, beta and exo genes (the bacteriophage *λ* red genes) under the control of an Arabinose-inducible promoter, and pSUB11, carrying the x3FLAG::KmR cassette. The cassette in pSUB11 was PCR-amplified with primers (forward: 5’-GAA GCA GGA AGA TAT GAT GCG CCT GCT AAG AGG CGG CAA CGA CTA CAA AGA CCA TGA CG-3’ and reverse: 5’-GGG TTA TAA GCA GGA CGG GCA AGC CCA CGG TGT AAA CCC GCA TAT GAA TAT CCT CCT TAG-3’), the 5’ ends of which were designed to target the 3’end of the gene of interest, digested with DpnI at 37°C for one hour and, upon purification, used for subsequent electroporation. Similarly, AhpC::6xHis-TEV-3xFLAG::TetR and SipA::6xHis-TEV-3xFLAG::TetR constructs were produced by amplification of the plasmid pJet1.2-Hfq-HTF-TetR (82) with the primers AhpC-forward 5’-AAA GAA GGC GAA GCG ACT CTG GCT CCA TCC TTA GAC CTG GTC GGT AAA ATC CGC TCT GC TGG ATC CAT GGA G-3’ and AhpC-reverse 5’-GTG AGC AGG CGA CGC CAA CGC AGC TAT GGC GTG AAA GAC GAC GGA AAT TTA CGC GTG AGG GGA TCT TGA AG-3’) or SipA-forward 5’-CCT GGC GTG GAT CGG GTT ATT ACT ACC GTT GAT GGC TTG CAC ATG CAG CGT CGC TCT GCT GGA TCC ATG GAG-3’ and SipA-reverse 5’-TTT GAC TCT TGC TTC AAT ATC CAT ATT CAT CGC ATC TTT CCC GGT TAA TTA CGC GTG AGG GGA TCT TGA AG-3’). Prior to electroporation, PCR products were digested with Exo1 and DpnI during one hour at 37°C followed by ethanol precipitation and verification on Agarose gels.

*Salmonella* Typhimurium SL1344 harbouring the plasmid pKD46 was grown in LB containing Ampicillin (100 μg/ml) and L-Arabinose (100 mM) at 28-30°C to an OD600 of 0.8. Cells were incubated on ice for 15 min, centrifuged at 3220 x g for 5 min at 4°C and resuspended in ice-cold water. The wash was repeated three times. On the final wash, cells were resuspended in 300 μl water and electroporated with 200 ng of PCR product. Cells were recovered for one hour in LB at 37°C on a tabletop thermomixer at 600 rpm, plated on LB agar with Kanamycin (50 μg/ml) or Tetracyclin (100 μg/ml) overnight. The following day, 10 colonies per strain were picked, resuspended in PBS and streaked on plates containing Ampicillin or Kanamycin(or Tetracyclin) and incubated at 37-40°C. Colonies that showed resistance to Kanamycin (or Tetracyclin) but not to Ampicillin were selected for further analysis, and the correct expression of the epitope tag was verified by western blot.

### PTex (phenol-toluol extraction)

HEK293 suspensions in 600 μl of DPBS (5-8 × 10^6^ cells, +/−CL) were mixed with 200 μl of each: neutral phenol (Roti-Phenol, Roth 0038.3), toluol (Th.Geyer, 752.1000) and 1,3-bromochloropropane (BCP) (Merck, 8.01627.0250) for 1 min (21°C, 2.000 r.p.m, Eppendorf ThermoMixer) and centrifuged 20.000 × g 3 min, 4°C. The upper aqueous phase (Aq1) was carefully removed and transferred to a new 2 ml tube containing 300 μl of solution D (5.85 M guanidine isothiocyanate (Roth, 0017.3); 31.1 mM sodium citrate (Roth, 3580.3); 25.6 mM N-lauryosyl-sarcosine (PanReac AppliChem, A7402.0100); 1% 2-mercaptoethanol (Sigma)). Then, 600 μl neutral phenol and 200 μl BCP were added, mixed and centrifuged as before. After phase separation, the upper 3/4 of Aq2 and the lower 3/4 of Org2 were removed. The resulting interphase (Int2) was kept in the same tube and mixed with 400 μl water, 200 μl ethanol p.a., 400 μl neutral phenol and 200 μl BCP (1 min, 21°C, 2.000 r.p.m, Eppendorf ThermoMixer) and centrifuged as previously. Aq3 and Org3 were carefully removed, while Int3 was precipitated with 9 volumes of ethanol (−20°C, 30 min to overnight). Samples were centrifuged during 30 min at 20.000 × g, pellets dried under the hood for max. 10 min and solubilised with 30 μL Laemmli buffer at 95°C for 5 min; or the indicated buffer/temperature according to the downstream application. For global mapping of RNA-protein interactions in HEK293 cells, 3 replicates from each UV irradiation energy were used.

### Hot-PTex from mouse tissue

260 mg of mouse-brain tissue was disrupted by cryogenic grinding. Immediately after, 130 mg were irradiated with 0.75 J/cm^2^ UV light (λ= 254 nm) in a CL-1000 ultraviolet cross-linker device (Ultra-Violet Products Ltd). Samples (−/+CL) were resuspended in DPBS on ice. 600 μl aliquots (32.5 mg) were mixed during 5 min at 65°C in the presence of 0.5 g of low-binding zirconium beads (100 μ, OPS Diagnostics, LLC) and the phenol-toluol-BCP mix described above. After centrifuging (20.000 × g 3 min, 4°C) Aq1 was carefully removed. Consecutive extractions were done as described above with the solely difference that mixing steps were performed at 65°C (2.000 r.p.m, Eppendorf ThermoMixer).

### Hot-PTex from *Salmonella*

For global mapping of RNA-protein interactions in *Salmonella* cells from two biological replicates were used. Bacteria attached to filters obtained as described in the section “Bacterial cell culture and *in vivo* cross-linking” were collected with 12 ml of DPBS and aliquots of 4 ml were pelleted at 20.000 × g, 2 min, 4°C. A modification of the step 1 was introduced in HOT-PTex in order to improve removal of free-proteins, as follows: bacterial pellets (−/+CL) resuspended in 400 μl of DPBS supplemented with EDTA (5mM) were mixed with phenol-toluol-BCP (200 μl each), and 0.5g of zirconium beads during 5 min at 65 °C (2.000 r.p.m, Eppendorf ThermoMixer). After centrifugation at 20.000 × g, 3 min, 4°C, the upper aqueous phase (Aq1) was mixed again with the same volumes of phenol-toluol-BCP (without beads) during 1 min at 65°C. Then the aqueous phase was carefully transferred to a third tube where steps 2 and 3 of the PTex protocol were performed at 65°C. Detailed step-by-step protocols of PTex and Hot-PTex are available as Supplementary Methods.

### Analysis of individual PTex steps

PTex extractions were carried as described before, using sets of three tubes containing synthetic 30-50 nt RNAs 5′-labelled 32P-ATP, 200 ng pUC19 LacZ-containing fragment (817 bp, generated by DrdI, NEB) or 2-3×106 HEK293 cells spiked-in with 0.25 μg of Sxl-RBD4 per tube. PCRs were performed using primers designed to amplify endogenous chromosomal DNA, il-3 gene (574 bp, forward primer: 5′-GAT CGG ATC CTA ATA CGA CTC ACT ATA GGC GAC ATC CAA TCC ATA TCA AGG A-3′ and reverse primer: 5′-GAT CAA GCT TGT TCA GAG TCT AGT TTA TTC TCA CAC-3′), or the LacZ gene present in the pUC19 linear fragment (324 nt, forward 5′- AGA GCA GAT TGT ACT GAG-3′and M13-reverse 5′-CAG GAA ACA GCT ATG ACC). DNA and RNA samples were electrophoresed in Agarose 1.0-1.5% or TBE-Urea PAGE 12%, respectively. Radioactivity was detected by phosphoimaging, while Sxl-RBD4, endogenous HuR or ACTB in HEK293/Sxl-RBD4 spiked-in proteins were analysed by Western blot with specific antibodies as described in the “Western Blotting” section.

### RNaseA Digestion and Electrophoretic Mobility Assay

Crosslinked and non-cross-linked HEK293 cell suspensions were subjected to PTex, the resulting pellets were solubilised in 150 μl of buffer TED (20 mM Tris, 1 mM EDTA, 0.03% DDM [n-Dodecyl β-D-maltoside]) at 56°C during 20 min, followed by incubation at 75°C during 20 min. Samples were mixed with RNaseA (2 ng) and incubated at 37°C; aliquots of 20 μL were taken at different time points: 0, 1, 5, 10, 30 and 60 min, immediately mixed with 5 μL of 6x Laemmli buffer, heated at 95°C for 5 min and used for SDS-PAGE.

### RNase treatment prior PTex

For PTex extractions shown in Fig. 1g, suspensions of 2-3×10^6^ HEK293 cells/ml (−/+ CL) were treated with 2000 U/ml benzonase (Merck, 70664) in the recommended buffer (50 mM Tris, 1 mM MgCL2), pH 8.0) during one hour at 37°C and 1000 r.p.m (ThermoMixer, Eppendorf). Untreated cells (−/+CL) were used as controls. PTex extraction were performed as described in “PTex (phenol-toluol extraction)”. After ethanol precipitation, pellets were directly solubilised in 40 μl Laemmli buffer (2×). SDS-PAGE and Western blots were performed as indicated below.

### *In-vitro* transcription of Sxl-RBD4 target RNAs

The T7 promoter and a sxl-target DNA sequence 5′-GAT CCG GTC ATA GGT GTA AAA AAA GTC TCC ATT CCT ATA GTG AGT CGT ATT AA-3′ was cloned into pUC19 using the restriction enzymes BamHI and HindIII. The resulting plasmid was named pUC19-sxl-target. Templates for RNAs of 87 and 191 nt length were generated using DNA restriction fragments from the pUC19-sxl-target plasmid (HindIII+EcoRI - 87 bp; HindIII+PvuI - 191 bp). The 30 nt RNA was synthesised as described before (83) by hybridising two complementary sequences containing the T7 RNA polymerase promoter 5′-TAA TAC GAC TCA CTA TAG-3′ and the template sequence 5′-GGT CAT AGG TGT AAA AAA ACT CTC CAT TCC TAT AGT GAG TCG TAT TAA-3′, followed by T7 run-off transcription. T7 RNA polymerase and restriction enzymes were purchased from New England Biolabs. Plasmids were purified using the NucleoBond Xtra Midi kit (Macherey-Nagel, 740410.100), DNA fragments by NucleoSpin Gel and PCR clean-up (Macherey-Nagel, 740609.50) and RNA by acidic phenol extraction (18). RNA 5’-GAG UUU UUU UAC A-3’ (13 nt) was synthesised by Biomers (Ulm, Germany).

### *In vitro* cross-linking assays

40 μg of Sxl-RBD4 in 100 μl cross-linking buffer (CLB: 10 mM Tris pH 7.4, 50 mM KCl, 1 mM EDTA, 1 mM DDT) were mixed with *in vitro*-transcribed RNA (13, 30, 87 and 191 nt, 1.7-10 μM) harbouring one copy of the target motif 5’-GAG UUU UUU UAC A-3’, incubated at 4°C for 30 min and cross-linked with 0.25 J/cm^2^ of UV-254 nm, on ice. Afterwards, 98% of each sample was used for PTex extraction, while 2% of the sample were kept as input control for SDS-PAGE and western blotting to detect Sxl-RBD4.

### Quantification of PTex

In order to unbiasedly determine the overall yield of PTex, a quantification of the total amount of clRNPs from the cell suspension used as input is required, which is so fartechnically impossible. As an alternative, we prepared mRNA interactome capture (RIC) samples from +CL HEK293(1) to serve as a “only-clRNPs” starting material. RIC samples from five biological replicates were used for PTex extraction. After ethanol precipitation, PTex samples were washed once with 5 ml of cold ethanol to remove traces of phenol which could interfere with the quantification, and resuspended in 20-50 μl of water. The absorbance of RIC and PTex samples were measured at λ280 nm and λ260/280 nm in a Nanodrop 2000. Additionally, 2% and 45% of RIC and PTex samples, repectively, were digested with RNaseA (0.1 μg/μl) at 37°C during 40 min. Intact and digested samples were loaded in Bis-Tris-MOPS gels 4-12% (NuPage, Invitrogen), transferred to nitrocellulose membranes and blotted to detect HuR (see below). A selection of western blot images generated during this study (Fig. S13,S14) were used to calculate the performance of PTex in terms of yield and specific-clRNP enrichment by densitometry analysis using ImageJ(84).

### Western blotting

Western blotting was performed following standard techniques. Samples were electrophoresed on SDS-PAGE gradient gels 4-20% (TGX stain free, Bio-Rad) or Bis-Tris 4-12% (NuPAGE, Invitrogen) and proteins transferred onto nitrocellulose membranes 0.2 μ (Bio-Rad). Membranes were blocked during 30 min with PBST-M (10 mM phosphate, 2.7 mM potassium chloride, 137 mM sodium chloride, pH 7.4 0.1% tween 20 (Sigma), 5% milk) and incubated with 0.1-1.0 μg/ml of the respective antibody overnight at 4° C (or 2h room temperature). Primary antibodies targeted the proteins HuR (proteintech, 11910-1-AP), ABCF2 (proteintech, 10226-1-AP), CCT7 (15994-1-AP) hnRNPL (proteintech, 18354-1-AP), FUS (abcam, ab124923), GAPDH (proteintech, 10494-1-AP), alpha-enolase (ENO1, proteintech,11204-1-AP), PTBP1 (abcam, ab133734), PABPC1 (proteintech, 10970-1-AP), ACTB (proteintech, 66009-1-Ig), Histone H3 (abcam, ab21054), FLAG-tag (Sigma, A8592), or Sxl-RBD4 (DHSB, anti-Sxl hybridome culture supernatant M114 1:20). Monoclonal mouse anti-Sxl antibodies (M18 and M114) were developed by P. Schedl and obtained from the Developmental Studies Hybridoma Bank (DHSB, created by the NICHD of the NIH and maintained at The University of Iowa, Department of Biology, Iowa City, IA52242). Antibody binding was detected using anti-mouseHRP (proteintech, SA00001-1), antimouse-AlexaFluor680 (Invitrogen, A32729) anti-rabbitHRP (proteintech, SA00001-2), or anti-rabbit-AlexaFluor488 (Invitrogen, A32731) and Clarity ECL Western Blotting Substrate for chemiluminescence in a ChemiDocMP imaging system (BioRad). All full blots are shown in Supplementary Figure S8.

### Immunoprecipitation and PNK assay

Immunoprecipitation of bacterial FLAG-tagged proteins and radioactive labeling of RNA by PNK was performed as described by (63).

### Protein purification

Recombinant Sxl-RBD4 protein (Sxl amino acids 122-301) was purified essentially as described before (22). In brief, after IPTG induction for 4h at 23°C in *E. coli* (BL21Star [Invitrogen] transformed with the Rosetta 2 plasmid [Merck]), cells were lysed in a buffer containing 20 mM Tris/Cl pH 7.5, 1M NaCl, 0.2 mM EDTA, 1 mM DTT, cOmplete Protease Inhibitor Cocktail [Roche] followed by centrifugation for 20min at 12,000 x g. The cleared lysate was then subjected to GSH-affinity chromatography using an ÄKTA FPLC system. Bound protein was eluted in a buffer containing 100 mM HEPES/KOH pH 8.0, 50 mM glutathione, 50 mM KCl, and 1 mM DTT. Fractions containing the protein were supplemented with 3C protease and dialyzed overnight against IEX buffer (20 mM HEPES/KOH pH 8.0, 50 mM KCl, 10% Glycerol, 0.2 mM EDTA, 0.01% NP-40) followed by ion exchange chromatography using a MonoS column. Fractions containing the pure protein were pooled, dialyzed against storage buffer (20 mM HEPES/KOH pH 8.0, 20% Glycerol, 0.2 mM EDTA, 0.01% NP-40, 1 mM DTT) and stored at 80°C.

### MS sample preparation

The mass spectrometry proteomics data (HEK293) have been deposited to the ProteomeXchange Consortium (http://proteomecentral.proteomexchange.org) via the PRIDE partner repository with the dataset identifier PXD009571. Human HEK293 and *Salmonella* cells were cultivated as described above and pellets used in PTex protocol. A minor fraction of initial input material was lysed and proteins denatured in 1% SDS and 0.1 M DTT Phosphate Buffer Solution (PBS) by boiling for 10 min at 95°C. After cooling, Benzonase was added for 30 min at 37°C before cell lysates were spun down and supernatants transferred to fresh tubes. Remaining input material was used for RBPs enrichment with PTex as described above. After PTex, RBPs were precipitated in 90% ethanol solution at −20°C and 30 minutes centrifugation at 20,000 × g at 4°C. Protein pellets were resuspended in 2 M urea in 50 mM ammonium bicarbonate (ABC) buffer and Benzonase was added for 30 min at 37°C to remove RNA. For silver staining, protein samples were directly loaded and separated on a pre-casted SDS-PAGE gel. Gels were fixed in 30% ethanol, 15% acetic acid solution in MiliQ water and incubated for 1 hour in 500 mM sodium acetate, 12 mM sodium thiosulfate, 0.125% glutaraldehyde, 25% ethanol solution. Gels were washed 3 times with MiliQ water for 10 minutes and stained with 0.1% Silver nitrate, 0.011% formaldehyde in MiliQ water solution for 30 minutes. Finally, gels were briefly rinsed with MiliQ water and reaction developed in 240 mM sodium carbonate, 0.01% Formaldehyde in MiliQ water solution. Reaction was stopped by addition of 50 mM EDTA.

For mass spectrometry analysis, proteins were precipitated with methanol-chloroform extraction (Wessel and Flugge 1984) and ressuspended in 8 M urea and 0.1 M Tris pH 8 solution. Proteins were reduced with 10 mM DTT at room temperature for 30 min and alkylated with 55 mM iodoacetamide at room temperature for 30 min in the dark. Proteins were first digested by lysyl endopeptidase (LysC) (Wako) at a LysC-to-protein ratio of 50:1 (w/w) at room temperature for 3 h. Afterwards, samples were diluted to 2 M final concentration of urea with 50 mM ammonium bicarbonate. Trypsin (Promega) digestion was performed at a trypsin-to-protein ratio of 50:1 (w/w) under constant agitation at room temperature for 16 h. Peptides were desalted with C18 Stage Tips (85) prior to LC-MS/MS analysis. Peptides were separated on a monolithic column (100 μm ID × 2,000 mm, MonoCap C18 High Resolution 2000 [GL Sciences] kindly provided by Dr. Yasushi Ishihama [Kyoto University]) using 6 hour gradient of increasing acetonitrile concentration at a flow rate of 300 nl/min, or on an in-house made C18 15cm microcolumns (75 μm ID packed with ReproSil-Pur C18-AQ 3-μm resin, Dr. Maisch GmbH) using 2 or 4 hours gradient of 5 to 50% increasing acetonitrile concentration at a flow rate of 200 nl/min. The Q Exactive instrument (Thermo Fisher Scientific) was operated in the data dependent mode with a full scan in the Orbitrap followed by top 10 MS/MS scans using higher-energy collision dissociation (HCD). All raw files were analyzed with MaxQuant software (v1.5.1.2) (86) using the label free quantification (LFQ) algorithm (87) with default parameters and match between runs option on. Database search was performed against the human reference proteome (UNIPROT version 2014-10, downloaded in October 2014) or the Salmonella Typhimurium reference proteome (UNIPROT version 2017, downloaded in August 2017) with common contaminants. False discovery rate (FDR) was set to 1% at peptide and protein levels.

### Bioinformatic analysis of PTex-purified proteins (HEK293)

The complete analysis is available as R notebook (Suppl_MS-HEK293_analysis). In short, we used LFQ MS intensities normalised to trypsin (which is constant in all samples). Potential contaminants, reverse and peptides only identified by modification were excluded from analysis. Fold changes were calculated by subtraction of the log2 values of LFQ intensity for proteins from UV cross-linked samples and non-cross-linked samples. Only proteins which were found in all replicates were processed further. Enrichment (CL/-CL) was calculated as described before (75): P-values were calculated from an Ebayes moderated t-test using the limma package (88) followed by Benjamini-Hochberg False Discovery Rate (FDR) correction. Only proteins with an adjusted p-value of 0.01 or smaller in all 3 cross-linking intensities were considered being enriched. GO analysis was performed using PANTHER V.11 (89). Domain enrichment was done using DAVID (90) searching the SMART (91) database.

### Bioinformatic analysis of PTex-purified proteins (Salmonella)

The complete analysis is available as R notebook (Suppl_MS-Salmonella_analysis). We used iBAQ-normalised values for the *Salmonella* analysis. Potential contaminants, reverse and peptides only identified by modification were excluded from analysis. Fold changes were calculated by subtraction of the log2 values of iBAQ intensity for proteins from UV cross-linked samples and non-cross-linked samples. Only proteins which were found in both replicates were taken into account (258 proteins; 172 with a log2 fold-change >0). Domain and GO terms were analysed using DAVID (90).

### RNASeq sample preparation

Cross-linked PTex samples and non-cross-linked control (CL-, 0.015, 0.15, 1.5 J/cm^2^) were proteinase K digested (1 h, 56°C) and the RNA recovered by acidic phenol extraction (18) using phase lock gel tubes (5Prime, 2302830). Libraries were created according to the “TruSeq Stranded Total RNA LT” protocol (Ilumina, 15032612) with the modification that we skipped the rRNA depletion step. We used adapters AR002,4,5-7,12,13-16,18,19. DNA concentration was determined by Qubit 3.0 Fluorometer (Life Technologies) and the quality of libraries assessed by a Bioanalyzer 2100, DNA 1000 Chip Kit (Agilent, 5067-1504). Sequencing was performed on a Illumina HiSeq4000. Note that we sequenced each condition in triplicates with the exception of input 0.015 J/cm^2^ for which we have duplicates. PTex RNA-seq data have been submitted to the NCBI Gene Expression Omnibus (GEO; http://www.ncbi.nlm.nih.gov/geo/) under accession number GSE113655. Read count data for RNA classes (Supplementary Fig. S20 are available in Supplementary Table S5).

### Bioinformatic analysis of PTex-purified RNA

RNASeq data were quality controled using fastqc (v0.11.2). We then mapped obtained reads against a single copies of human rRNA and tRNA sequences using bowtie2 (v2.2.6):

~~~
bowtie2 -no-unal -un
$LIBRARY_rRNA_not_aligned_reads.fastq
-al $LIBRARY_rRNA_aligned_reads.fastq
-x rRNA_db -U, $LIBRARY.fastq
> $LIBRARY_rRNA.sam 2»,
$LIBRARY_rRNA_alignment_stats.txt
~~~

We then used the remaining reads and mapped reads to the human genome (GRCh38.p12) and the corresponding comprehensive gene annotation file (gencode.v28.chr_patch_hapl_scaff.annotation.gtf) from GENCODE using the STAR aligner (v020201):

~~~
STAR -genomeDir PATH/TO/index
-readFilesIn $LIBRARY_rRNA_not_aligned_
reads.fastq -quantMode GeneCounts
~~~

After aligning the remaining reads with STAR, samtools calmd was used to annotate each read in the bam files with mutation information. Using the SAM.py parser from the pyCRAC suite (35), chromosomal locations of substitutions and deletions were extracted and counted. Only mutations that were unique to the UV-treated samples were considered. To normalize the data for sequencing depth, for each dataset the counts for substitutions and deletions were divided by the total number of mapped nucleotides, which provided an indication of mutation frequencies. To map the distribution of deletions and substitutions around AUG and Stop codons, CDS coordinates from the gencode.v28.chr_patch_hapl_scaff.annotation.gtf annotation files were extracted. Tables containing counts and chromosomal positions for each substitution and deletion were converted into gene transfer format (GTF) files using the GTF2 and NGSFormatWriters classes from the pyCRAC package. Subsequently, pyBinCollector from the pyCRAC package was used to map the distribution of substitutions and deletions around start and stop codons of protein-coding genes:

~~~
pyBinCollector.py -f mutations.gtf -gtf, annotationfile_CDS_coordinates.gtf -s 5end -a protein_coding -normalize -v -o dist_around_AUG.txt
pyBinCollector.py -f mutations.gtf -gtf, annotationfile_CDS_coordinates.gtf -s 3end -a protein_coding-normalize-v-o dist_around_STOP.txt
~~~

For the feature counts, count tables generated by STAR were used in conjunction with the gencode.v28.chr_patch_hapl_scaff.annotation.gtf annotation file.

### PAR-CLIP and pCLIP

We performed the PAR-CLIP protocol as described by (11, 27): HEK293 cells stably expressing FLAG-tagged HuR (ELAVL1), were grown until 90% confluence. The last 16 h of incubation, 200 mM 4SU was added. Living cells were irradiated with 0.15 J/cm^2^ 365 nm UV light, snap-frozen on dry ice and stored at −80°C until use. Cells were collected on different days, representing biological replicates. Cells (1.2 × 108 cells/replicate) were lysed on ice for 10 min with 3 ml lysis buffer (50 mM Tris-Cl pH 7.5 (Life Tech., 15567027), 100 mM NaCl (Life Tech. AM9760G), 1% (v/v) Nonidet P40 substitute (Sigma 74385), 0.5% (v/v) Sodium deoxycholate (AppliChem A1531) containing 0.04 U/ml RNasin (Promega, N2515) and 2x Complete Protease Inhibitor (Roche, 11697498001) and centrifuged 20,000 × g, 10 min, 4°C. Cleared lysates (1.5 mL/replicate) were digested with 8 U/mL TURBO DNase (ThermoFisher, AM2238) and 2 U/μL RNase I (ThermoFisher, AM2294) at 37°C for 4 min (replicate 1) or 3 min 15 sec (replicates 2 and 3). FLAG-tagged HuR was immunoprecipitated with 10 μg of antiFLAG monoclonal antibody (Sigma, F1804) bound to 40 μl of Protein G Dynabeads (Life Tech, 10004D). After extensive washes with high-salt buffer (50mM Tris-HCL pH 7.5, 1mM EDTA, 1% Nonidet P40, 0.1% (v/v) SDS, 1 M NaCl) beads were incubated with 1 U/μl of T4 polynucleotide kinase (T4PNK, NEB) and 0.5 μCi/μL of ^32^P-ATP. After radiolabelling, samples were splitted into 4 tubes and underwent three versions of CLIP:

### PAR-CLIP classic

Briefly, clHuR-RNA complexes were resolved by 4-12% Nu-PAGE MOPS (invitrogen) transferred to nitrocellulose and excised at a defined size-range (50 to 60 kDa). Proteins were digested from the membrane with proteinase K and the RNA recovered by acidic phenol/chloroform extraction and ethanol precipitation. Resulting RNA was ligated with 3’ adapter (5’App-NNN NTG GAA TTC TCG GGT GCC AAG G-3’InvdT) gel-slice isolated, the, ligated with the 5′adapter (5’-GUU CAG AGU UCU ACA GUC CGA CGA UCN NNN-3’) and purified again from PAGE-Urea gels by elution and ethanol precipitation (11, 27).

### PAR-CLIP on-beads

As an alternative, the ligation of the 3′/5′adapters can be achieved directly on the beads used in the affinity capture of the selected clRNP (28, 29). On-beads adapters ligation was done by incubating the FLAGclHuR-RNA beads with the 3′adapter in the presence of Rnl2(1–249)K227Q ligase and PEG-8000 overnight at 4°C. After washes, the 5′adapter was ligated using the Rnl1 enzyme, 2 h at 37°C as described in (29). Followed by protein electrophoresis (4-12% Nu-PAGE MOPS, Invitrogen), blotting to nitrocellulose membranes and band excision at the defined size-range (50 to 60 kDa). Proteins were digested from the membrane with proteinase K. RNA was then recovered by acidic phenol/chloroform extraction and ethanol precipitation.

### pCLIP

beads capturing clHuR-RNA complexes were subjected to 3′/ 5′adapters ligation as above. Immediately after the 5′adapter ligation beads were magnetically captured and resuspended in 600 μl solution D. After a short denaturation (95°C during 10 min), beads were separated with a magnet and the clHuR-RNA complexes recovered in the supernatant were subjected to the last two steps of the PTex protocol (20 min). PTex recovered interphases were precipitated with 9 volumes of ethanol on dry ice during 30 min, followed by 30 min centrifuging at 20.000 ×g, 4°C. Ethanol-precipitated pellets were digested with proteinase K and RNA isolated by phenol/chloroform extraction.

### Library preparation and RNAseq

PARCLIP RNA-seq data have been submitted to the NCBI Gene Expression Omnibus (GEO; http://www.ncbi.nlm.nih.gov/geo/) under accession number GSE113628. RNAs obtained from the three PARCLIP procedures (classic, on-beads, and pCLIP) were retrotranscribed into cDNA with the reverse transcription primer 5’-GCC TTG GCA CCC GAG AAT TCC A-3’ and the minimal PCR cycles were determined for each case. cDNA libraries were created by PCR using the forward primer 5’- AAT GAT ACG GCG ACC ACC GAG ATC TAC ACG TTC AGA GTT CTA CAG TCC GA-3’ and the Ilumina adapter index RPI 1-6, 8,10-11. Bands obtained at 150 bp were excised from 2 % agarose gels and purified using the Zymoclean Gel Recovery Kit (Zymo, D4002). DNA concentration and library quality was determined by Qubit Fluorometer dsDNA HS assay (Life Tech, Q32854) and BioAnalizer DNA HS Kit (Agilent 2100 Bioanalyzer; Agilent, 5067-4626). Libraries were sequenced in a NextSeq 500.

### CLIP data processing

The PAR-CLIP data was processed and annotated using the PARpipe pipeline (https://github.com/ohlerlab/PARpipe) around the PARCLIP data tailored peak caller PARalyzer (92) as described previously (26), with one modification. In brief, adapter sequences were trimmed retaining the four randomized adapter nucleotides on both read ends included during adapter ligation to serve as unique molecular identifiers during PCR-duplicate removal (read collapsing). Differences in the numbers of uniquely aligning reads were balanced by random subsampling prior to PARalyzer cluster identification. Annotation of identified binding sites (cluster) was simplified by grouping closely related sub-annotation categories (Supplementary Table S6).

### De novo motif discovery

For de novo motif finding we used Zagros (30) with default settings including RNA secondary structure information. As the majority of HuR cluster resided in intronic and 3’utr regions (70-80%, Supplementary Table S7) according to its reported functions, we use intronic and 3’utr cluster sequences as input. For DREME motif analysis, we used all PARalyzer-derived cluster sequences and ran DREME with default settings against shuffled background sequences allowing only sense strand motif search.

### Transcriptomic metacoverage

For depicting spatial preferences for mRNA binding, we selected genes previously used for RNA classification based on processing and turnover dynamics (93) (n=15120) present in GENCODE v19. To select transcripts, we ran RSEM (94) and retained transcripts with TPM >3. For each gene, we selected the transcript isoform with the highest isoform percentage or chose one randomly in case of ties (n=8298). The list of selected transcript isoforms was used to calculate the median 5’UTR, CDS and 3’UTR length proportions (5’UTR=0.06, CDS=0.53, 3’UTR=0.41) using R Bioconductor packages GenomicFeatures and GenomicRanges (95). For regions post annotated transcription ends and splice sites we chose windows of fixed sizes (TES 500 nt, 5’ and 3’ splice sites 250 nt each). We generated coverage tracks from the PARalyzer output alignment files and intersected those with the filtered transcripts. Each annotation category was binned according to its relative length and the coverage averaged within each bin. For intronic coverage, we averaged across all introns per gene, given a minimal intron length of 500 nt. All bins were stitched to one continuous track per transcript (n=6632 intron containing transcripts). Each library bam file was filtered to retain only PARalyzer cluster overlapping alignments. We required transcripts to have a minimal coverage maximum of >2. For each transcript we scaled the binned coverage dividing by its maximal coverage (min-to-1 scaling) to emphasize on spatial patterns independent from transcript expression levels. Next, we split transcript coverage in two parts, separating 5’UTR to TES regions and intronic regions. To generate the scaled meta coverage across all targeted transcripts per RBP, we used the heatMeta function from the Genomation package (96). For the 5’UTR to TES part, we scaled each RBP meta-coverage track independent of other libraries. For intronic sequences, we scaled each sample relative to all other sample. Finally, we clustered the meta-coverage tracks using ward.D clustering with euclidean distance.

### Genome browser vizualizations

PAR-CLIP alignments were visualized using Gviz (97).

## ACKNOWLEDGEMENTS

The authors wish to thank Florian Heyd and Marco Preussner for providing mouse brain samples, Jörg Vogel for providing Hfq-FLAG-tagged *Salmonella* and Stuart McKellar for help with cross-linking. We are grateful to Maja Köhn and Matthias Hentze for discussions and to Markus Landthaler for critically reading the manuscript. Work in the Beckmann lab is supported by the German Research Foundation (DFG; IRTG 2290 and ZUK 75/1 Project 0190-854599). Work in the Medenbach lab is supported by the DFG (SFB 960/2, B11), the Bavarian State Ministry for Education, Science and the Arts (Bavarian Research Network for Molecular Biosystems, BioSysNet), and the German Federal Ministry of Education and Research (BMBF, 01ZX1401D). The Granneman lab is funded by a Medical Research Council non-clinical Senior Research fellowship (MR/R008205/1). The Ohler lab is funded by NIH TRA R01 GM104962. ECU is supported by the Joachim Herz Foundation.

## AUTHOR CONTRIBUTIONS

E.C.U. and B.M.B designed the project; E.C.U., C.H.V-V., H.H.W, D.F., R.M performed the research; E.C.U., C.H.V-V., T.H., H.H.W, S.G, M.S. and analysed the data; E.C.U, S.G., U.O, J.M. and B.M.B wrote the paper with input from all authors.

