## Supplementary Information for "Purification of Cross-linked RNA-Protein Complexes by Phenol-Toluol Extraction"

#### Supplementary Figures

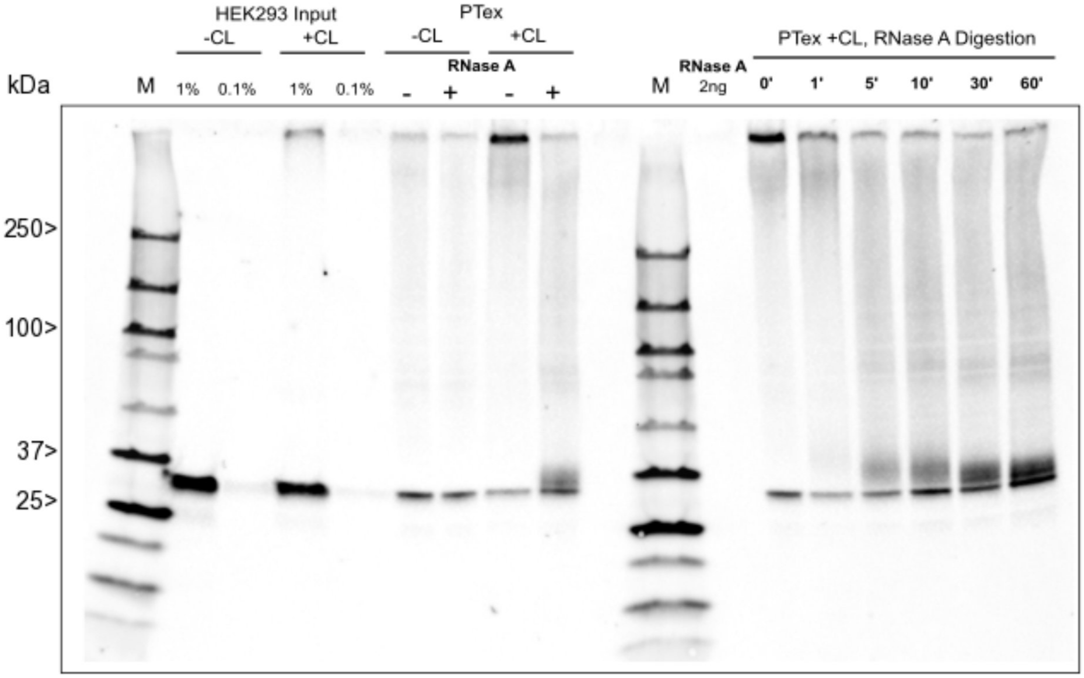


Figure S1: Western blot anti HuR (ELAVL1; 35 kDa) from HEK293 cells. Input: After UV cross-linking (+CL), a second band appears at the upper edge of the gel/at the gel pocket. To show that this signal is indeed caused by formation of RNA-protein cross-links, we performed PTex and incubated purified complexes (clHUR) with RNase A (0-30 min incubation time). The shifted complexes are depleted in a RNase- digestion-dependent fashion and a signal reconstitutes running at slightly higher MW than free HuR. Note that RNase digestion cannot completely remove cross-linked RNA, causing a slower migration in SDS-PAGE due to the additional mass of the leftover RNA.

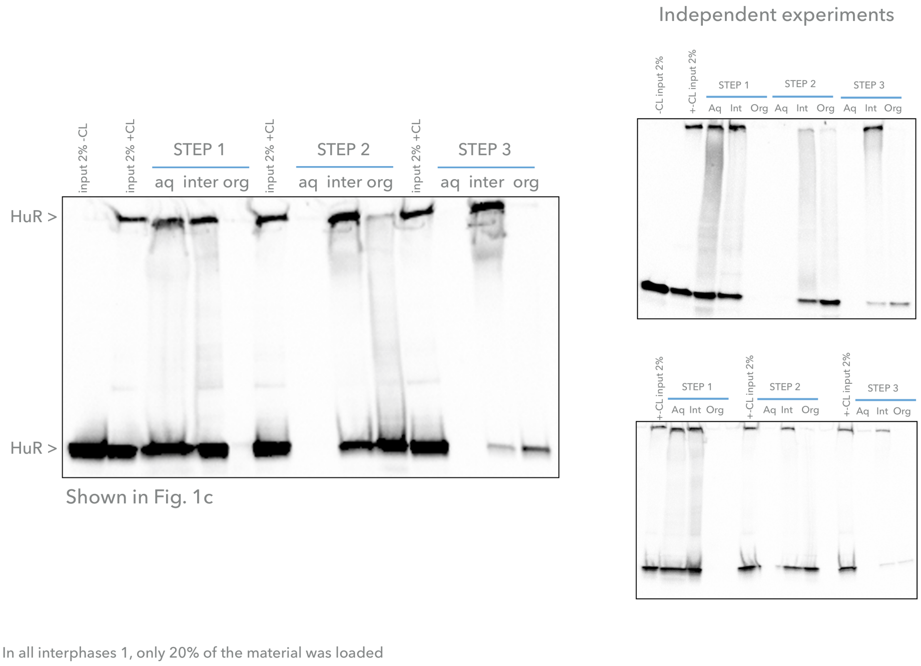


Figure S2: Full blots for Fig. 1c. Antibody against HuR.

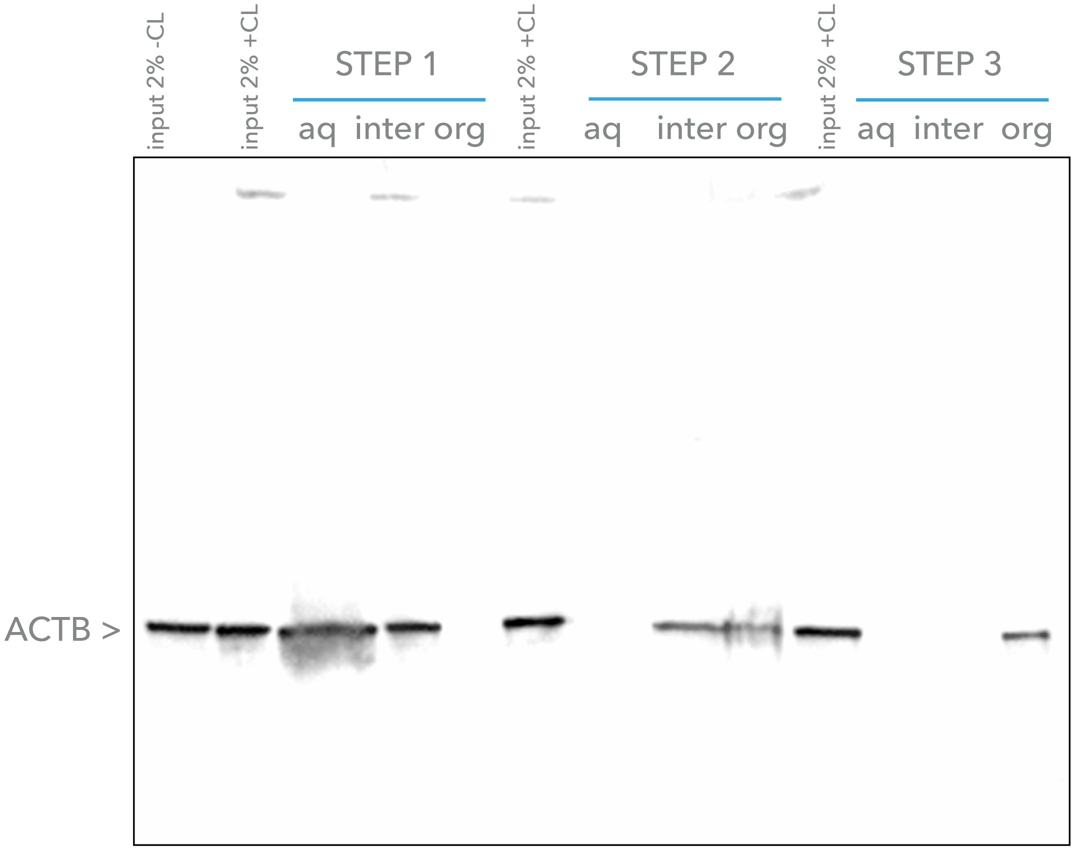


Figure 1C. PTex steps of HEK293 cells. Western blot against ACTB

Figure S3: Full blots for Fig. 1c. Antibody against beta-actin.

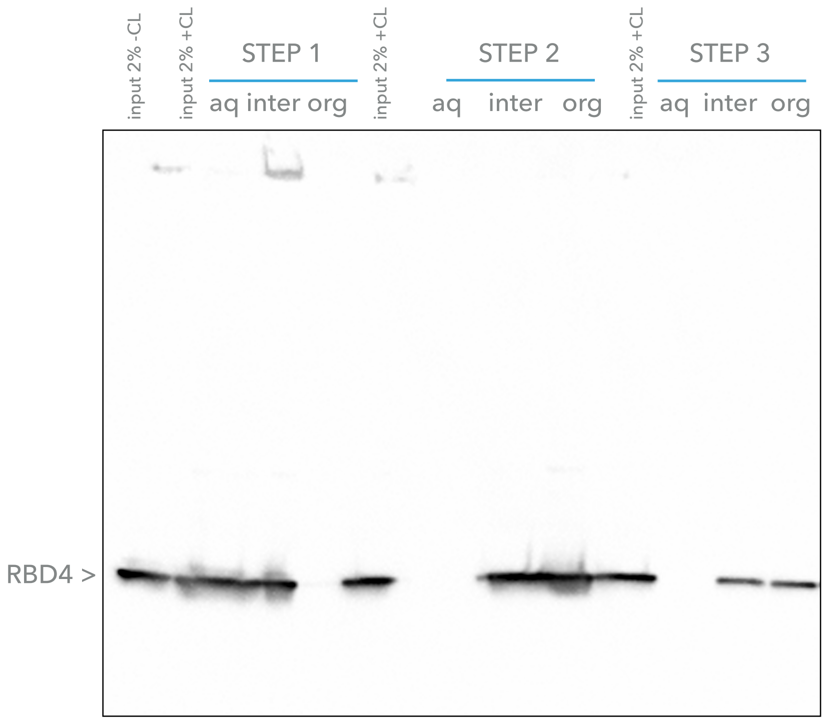


Figure 1C. PTex steps of HEK293 cells spiked-in DmRBD4. Western blot against RBD4

Figure S4: Full blots for Fig. 1c. Antibody against Sxl RBD4 spike-in protein.

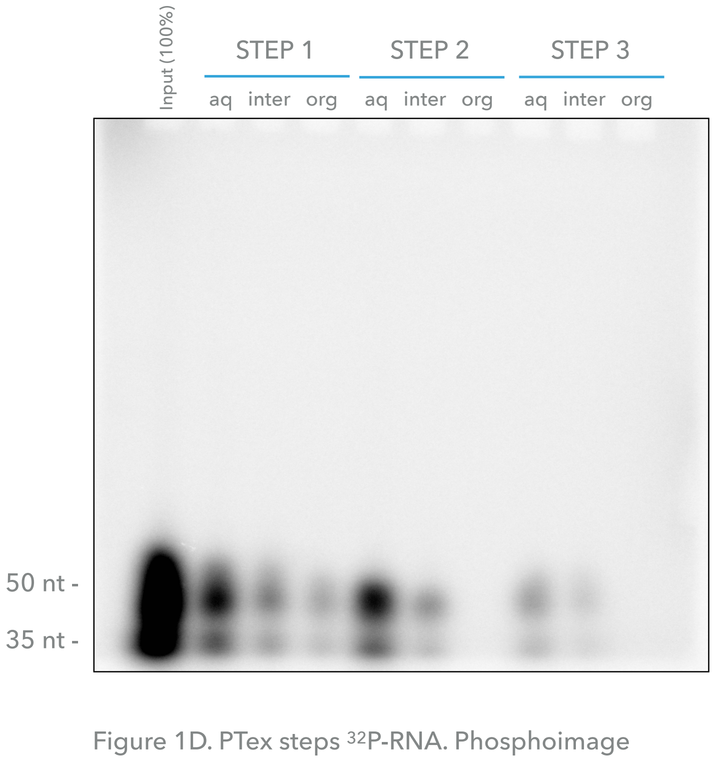


Figure S5: Full autoradiography for Fig. 1d.

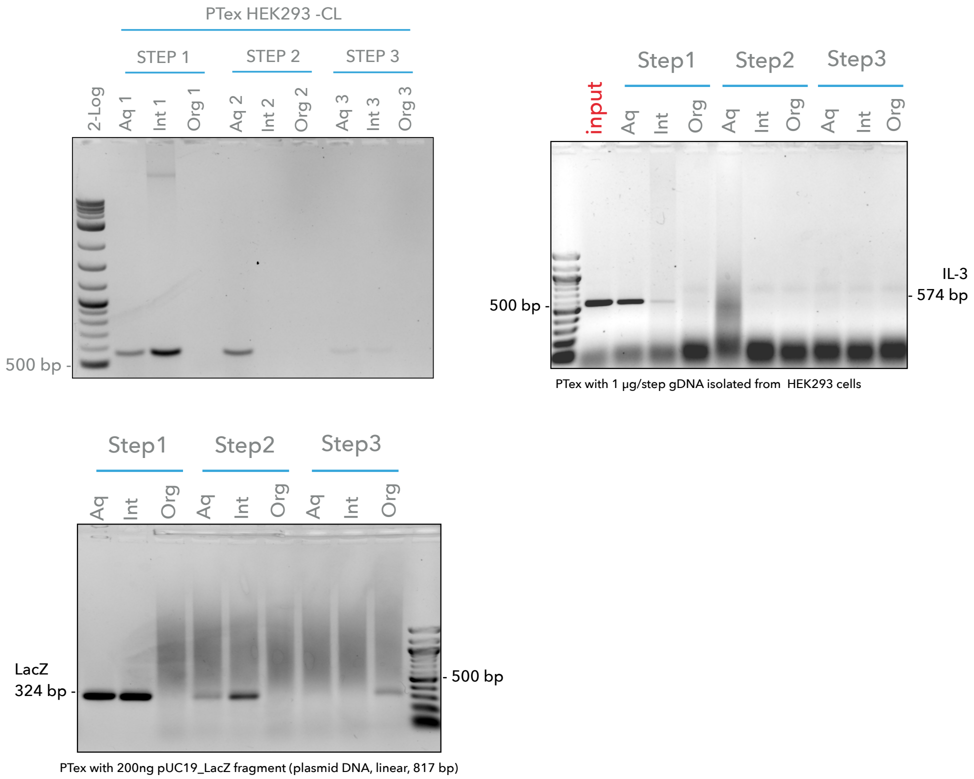


Figure S6: Full agarose gel images for Fig. 1e.

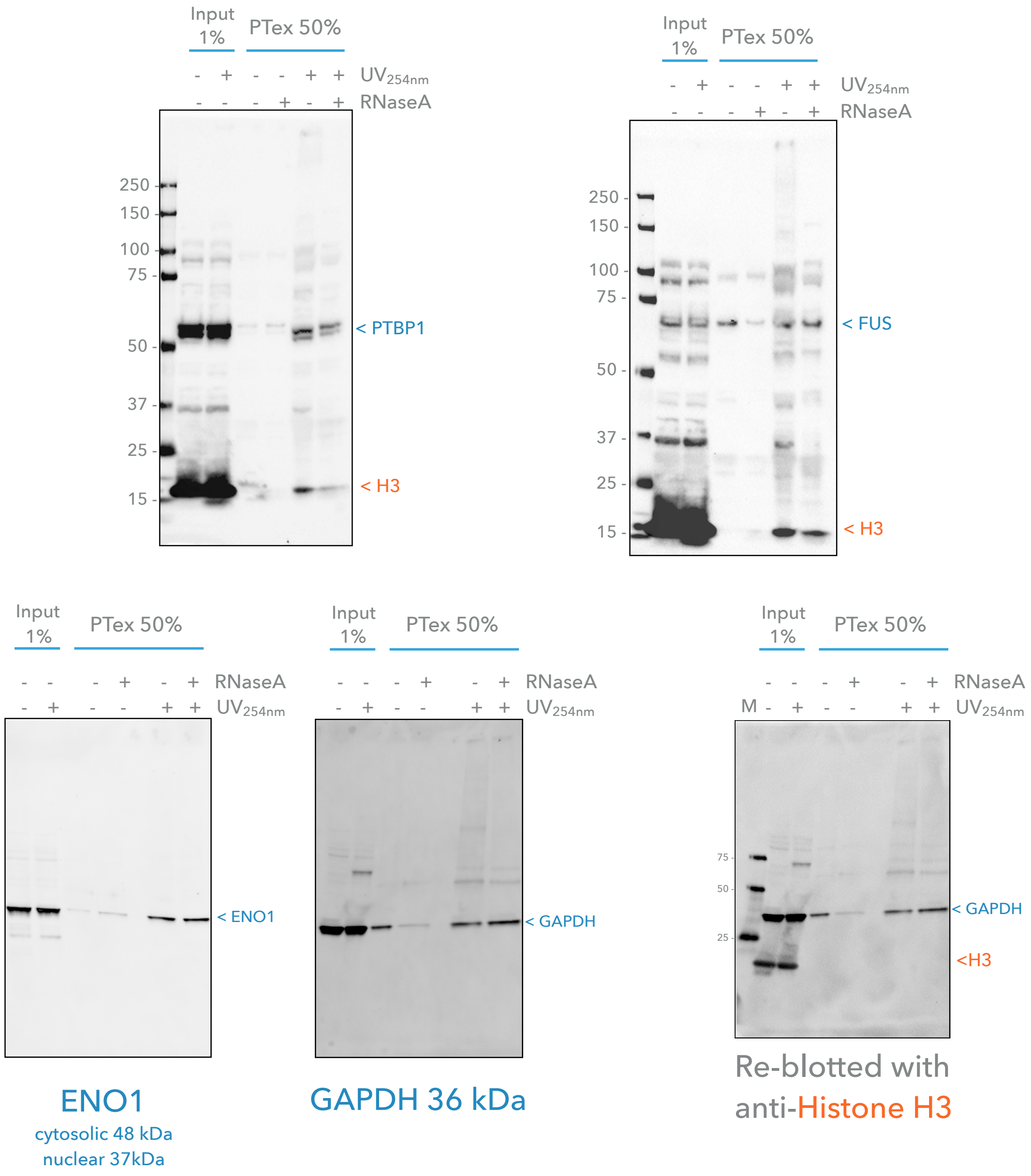


Figure S7: Full blots for Fig. 1f.

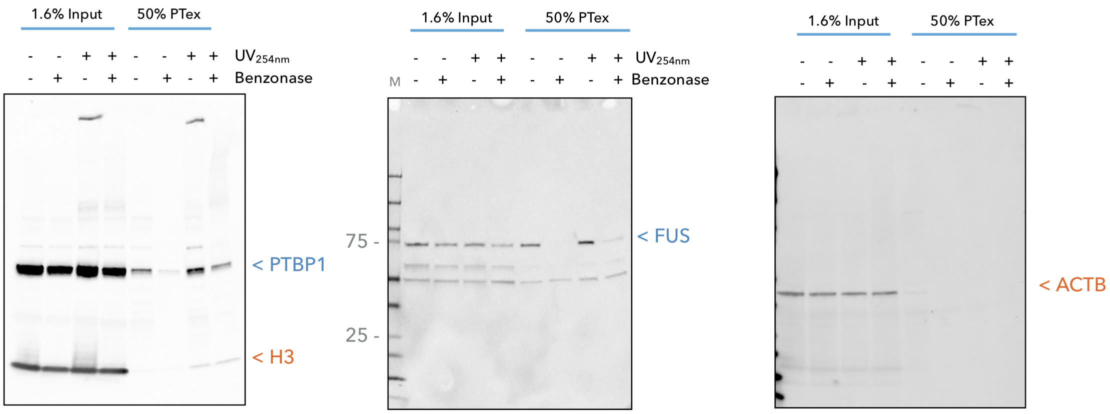


Figure S8: Full blots for Fig. 1g. HEK293 cellular suspensions (-/+ CL, 2-3x106 cells/mL) were incubated with 2000 U/ml of benzonase at 37C, 1h, 1000 r.p.m. (Thermomixer, Eppendorf). Non-treated samples were kept as -Benzonase controls. For PTex, 600 $\mu$l per condition were used.

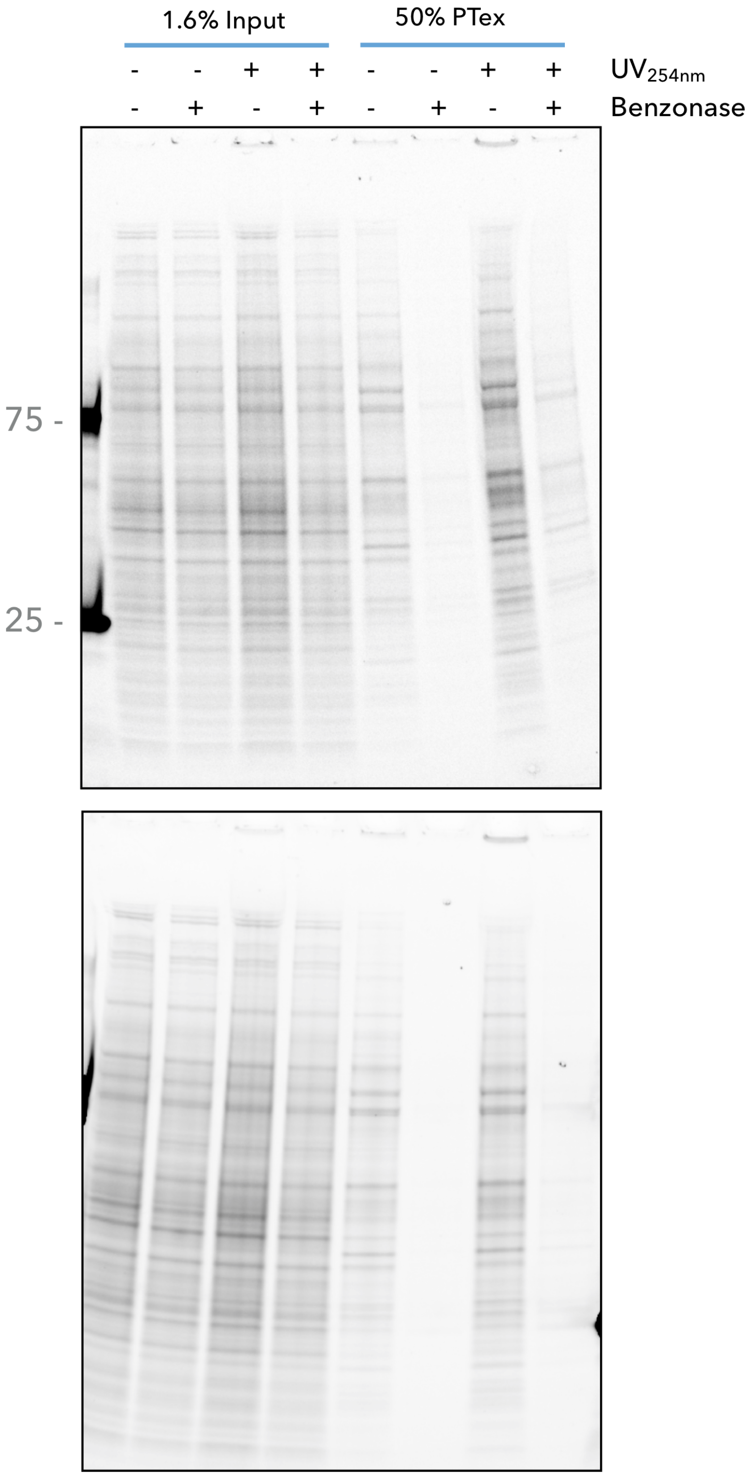


Figure S9: RNase (benzonase) treatment prior to PTex. HEK293 cellular suspensions (-/+ CL, 2-3x106 cells/mL) were incubated with 2000 U/ml of benzonase at 37C, 1h, 1000 r.p.m. (Thermomixer, Eppendorf). Non-treated samples were kept as -Benzonase controls. For PTex, 600 $\mu$l per condition were used. SDS-PAGE 4-20%, TGX Stain-Free, BioRad. Gels were UV activated during 2.5 min prior imaging (ChemiDoc).

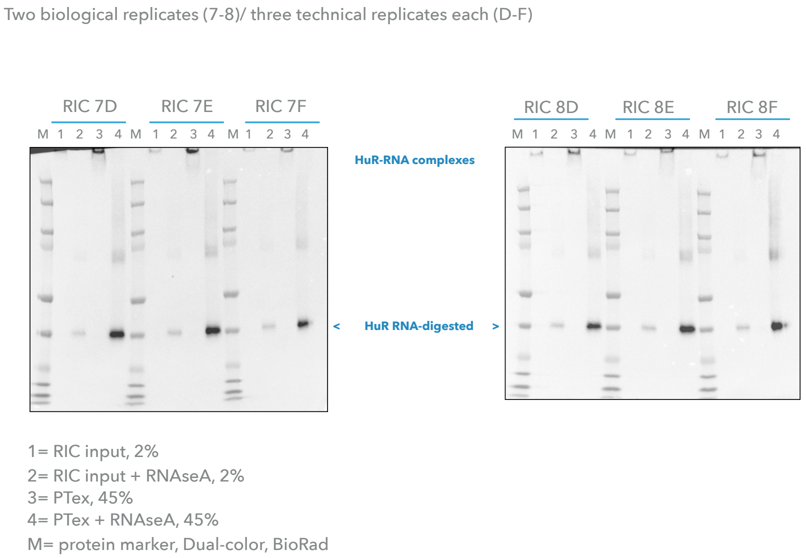


Figure S10: Full blots for Fig. 2a and 2b. Antibody staining against HuR. Note that RNase was added after RIC/PTex to shift cross-linked protein back to its original molecular mass.

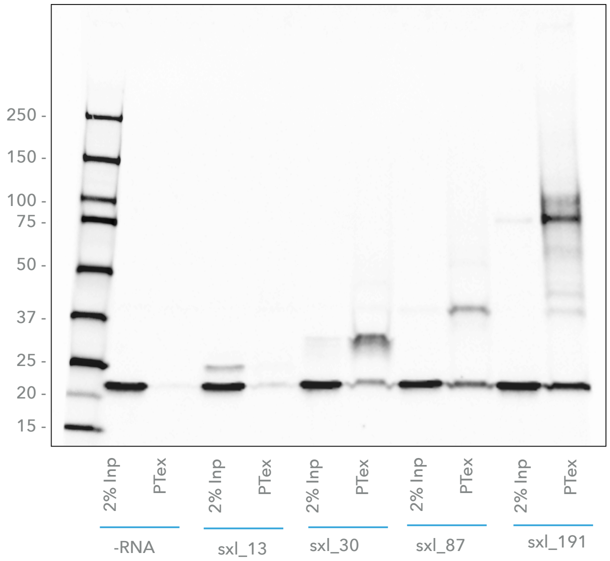


Figure S11: Full blot for Fig. 2c. Antibody staining against Sxl RBD4.

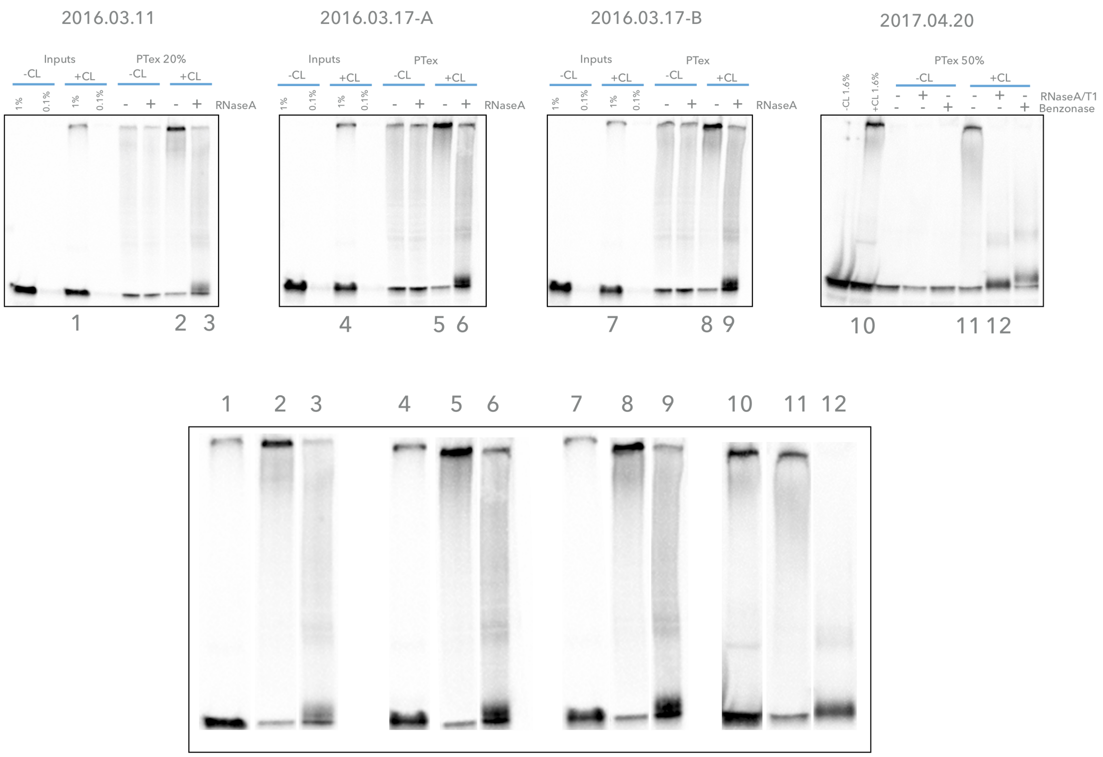


Figure S12: Western blots used for densitometry in Fig. 2f. Antibody staining against HuR.

ImageJ
https://imagej.nih.gov/ij/

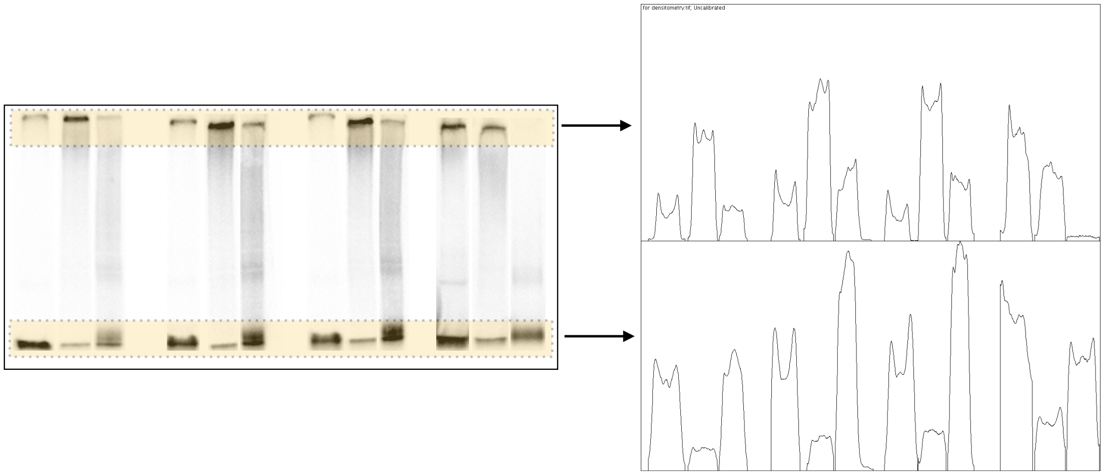


Figure S13: Blot areas used for densitometry in Fig. 2f. Antibody staining against HuR.

ImageJ
https://imagej.nih.gov/ij/

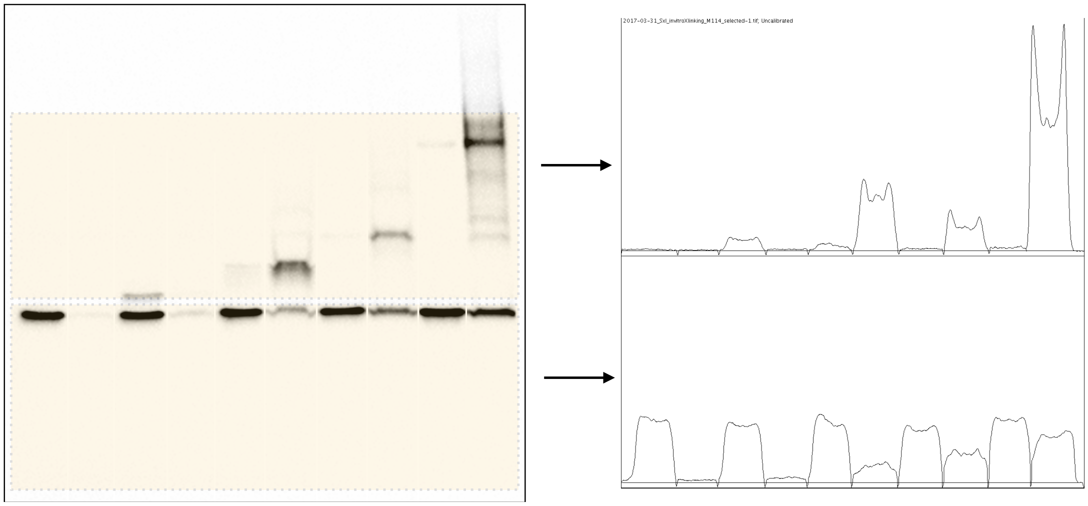


Figure S14: Blot areas used for densitometry in Fig. 2f. Antibody staining against Sxl RBD4.

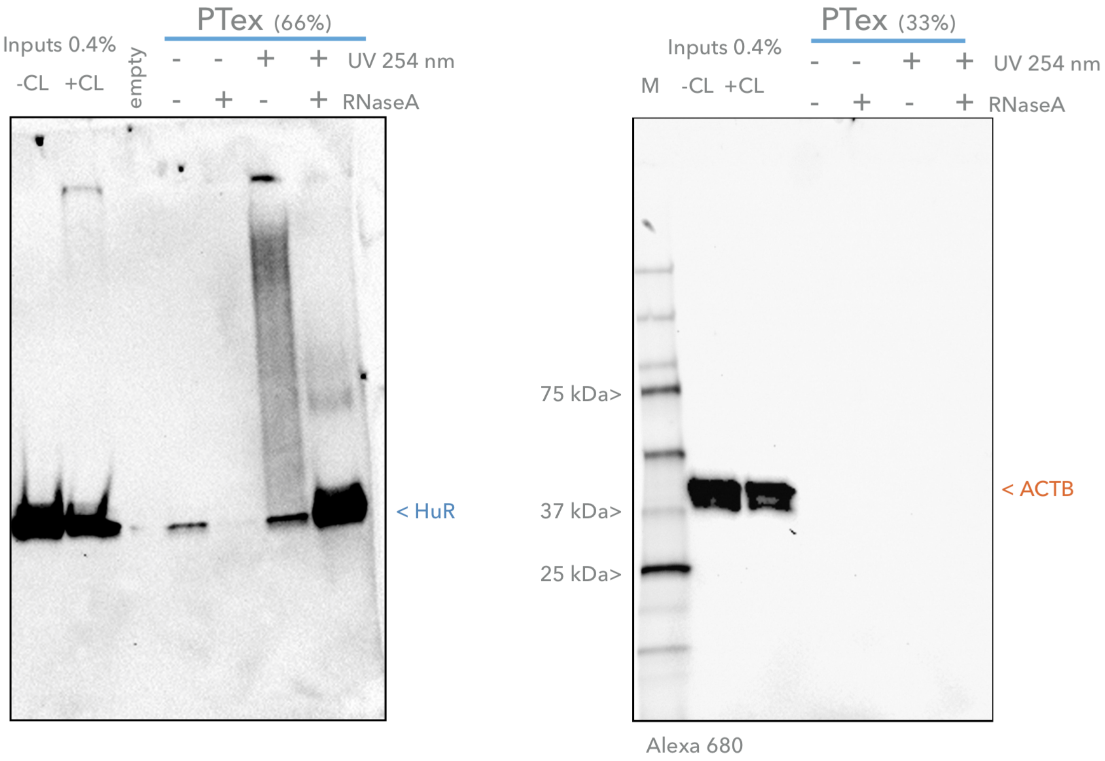


Figure S15: Full blots of PTex from mouse brain tissue (Fig. 3). Antibodies against HuR and beta-actin. RNase was added after PTex.

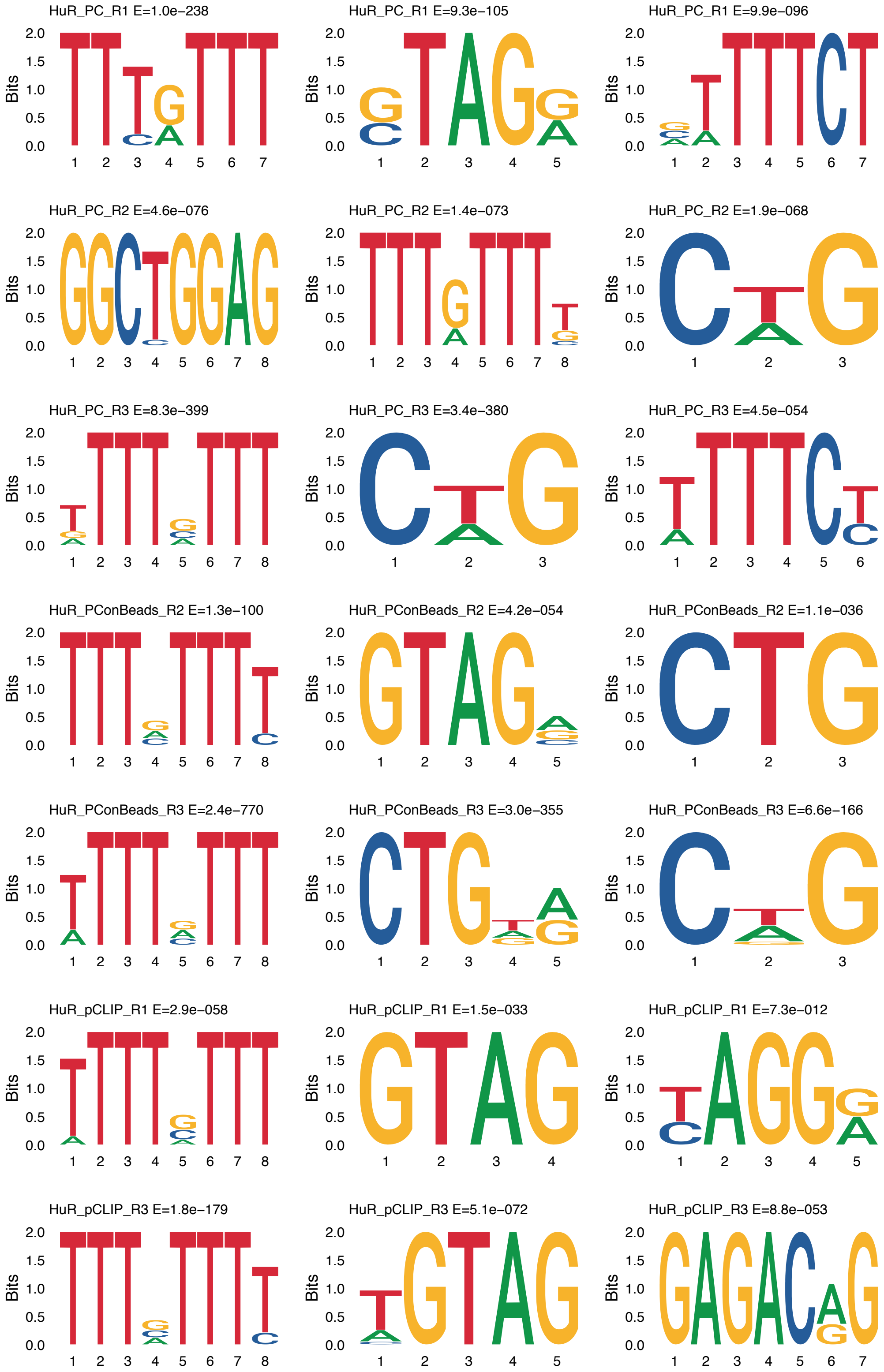


Figure S16: DREME motif analysis of HuR-bound RNA from PAR-CLIP/PAR-CLIP on beads/pCLIP (Fig.4).

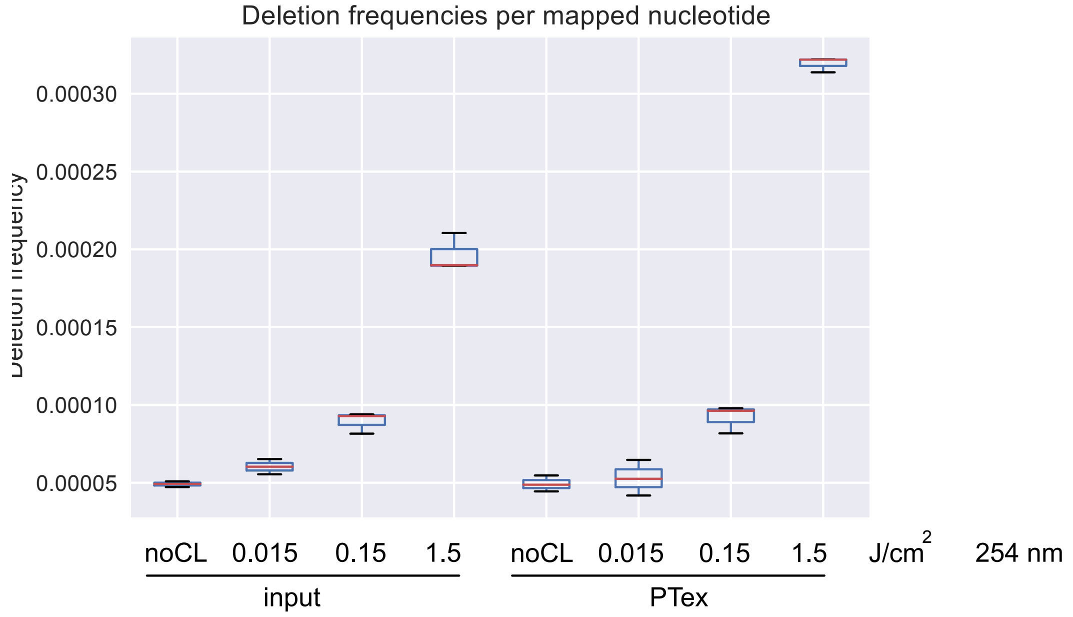


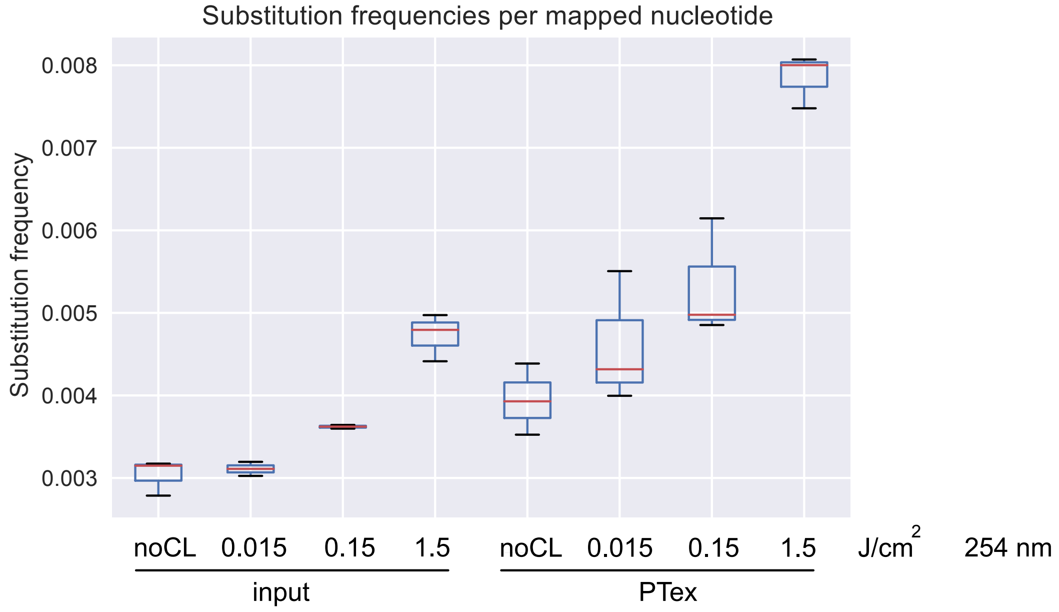


Figure S17: Deletion and Substitution frequencies for input and PTex-purified samples from HEK293 RNA. Corresponds to Fig. 5.

Deletions

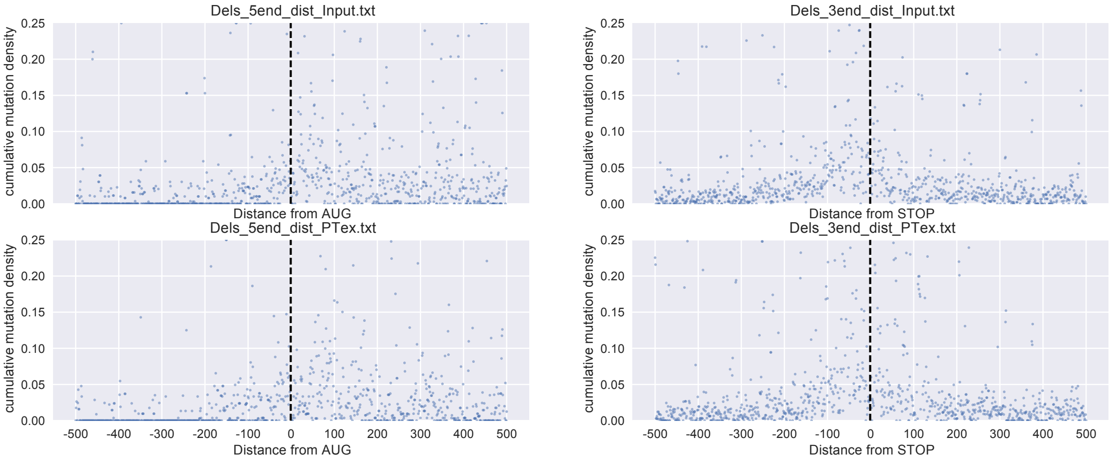


Substitutions

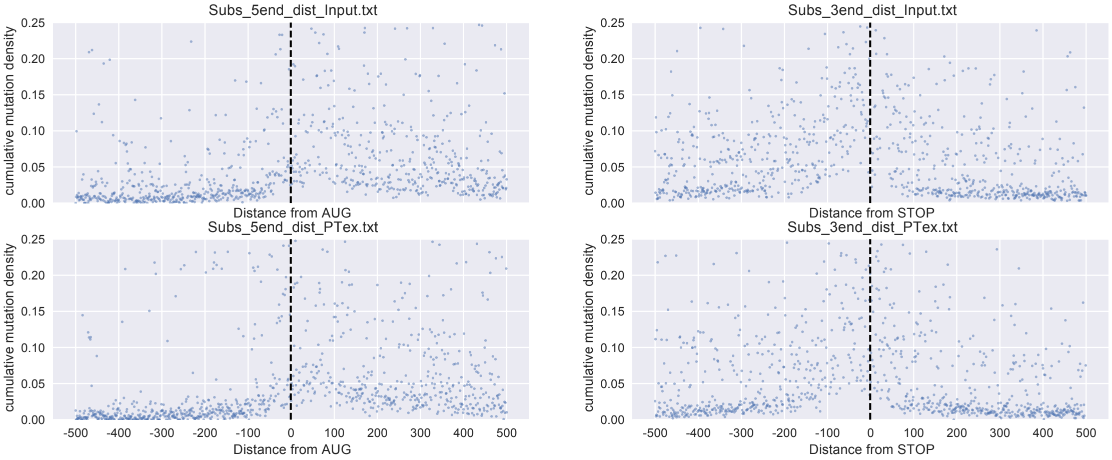


Figure S18: Mapping of deletions and substitutions in RNA transcripts from coding regions. Input (= total RNA) or PTex samples were analysed and mutations plotted relative to the AUG or STOP codon. Corresponds to Fig. 5.

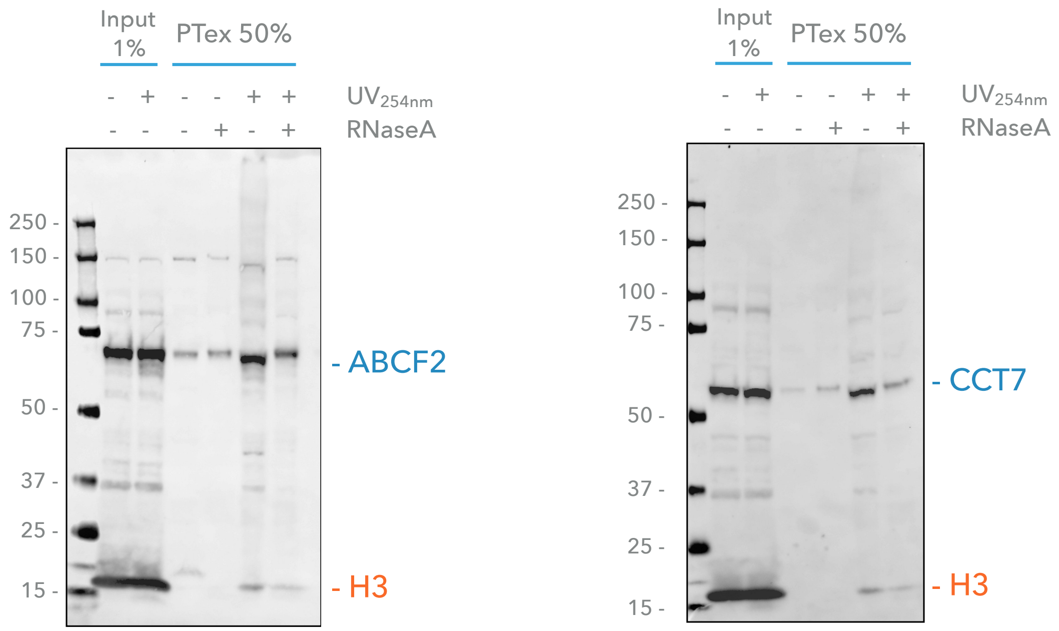


Figure S19: Full blots of PTex from PTex-enriched human proteins ABCF2 and CCT7 (Fig. 5).

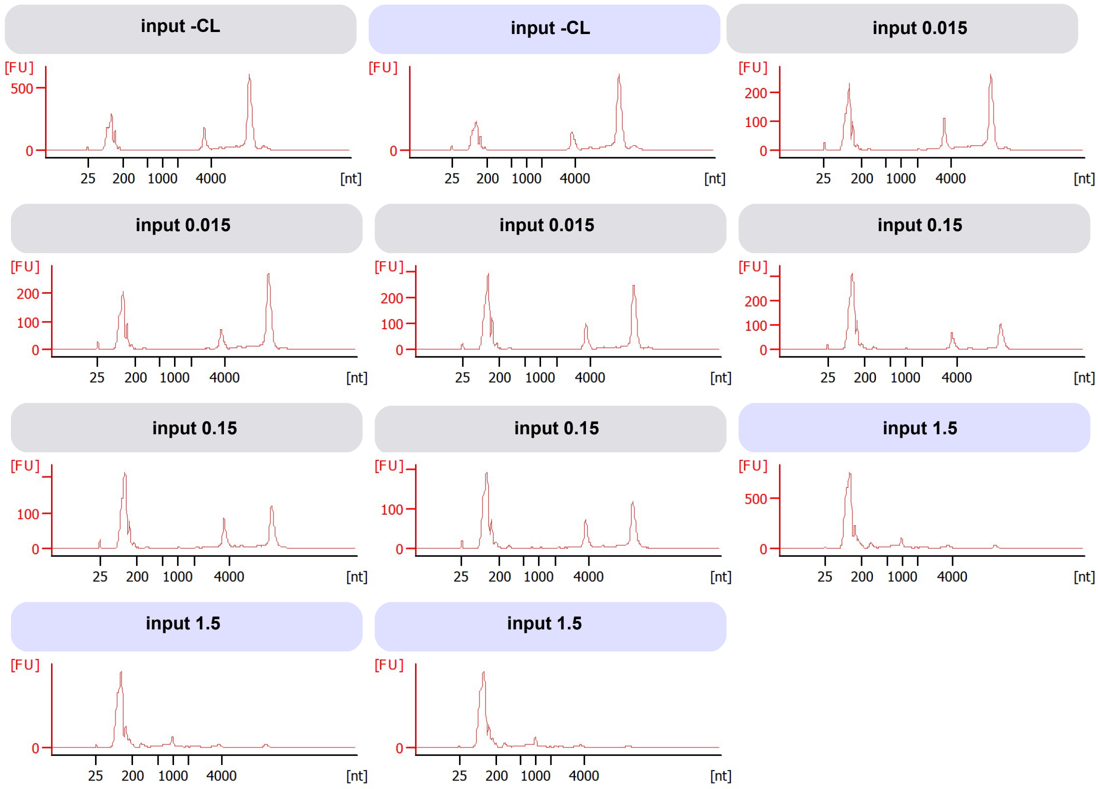


Figure S20: Agilent Bioanalyzer 2100 Total RNA Pico Chip analysis of total RNA preparations from HEK293 cells. Cells were not UV irradiated (-CL) or irradiated with 0.015, 0.15 or 1.5 J/cm2 254 nm wavelength before RNA extraction. The two ribosomal RNA peaks are indicative for RNA integrity.

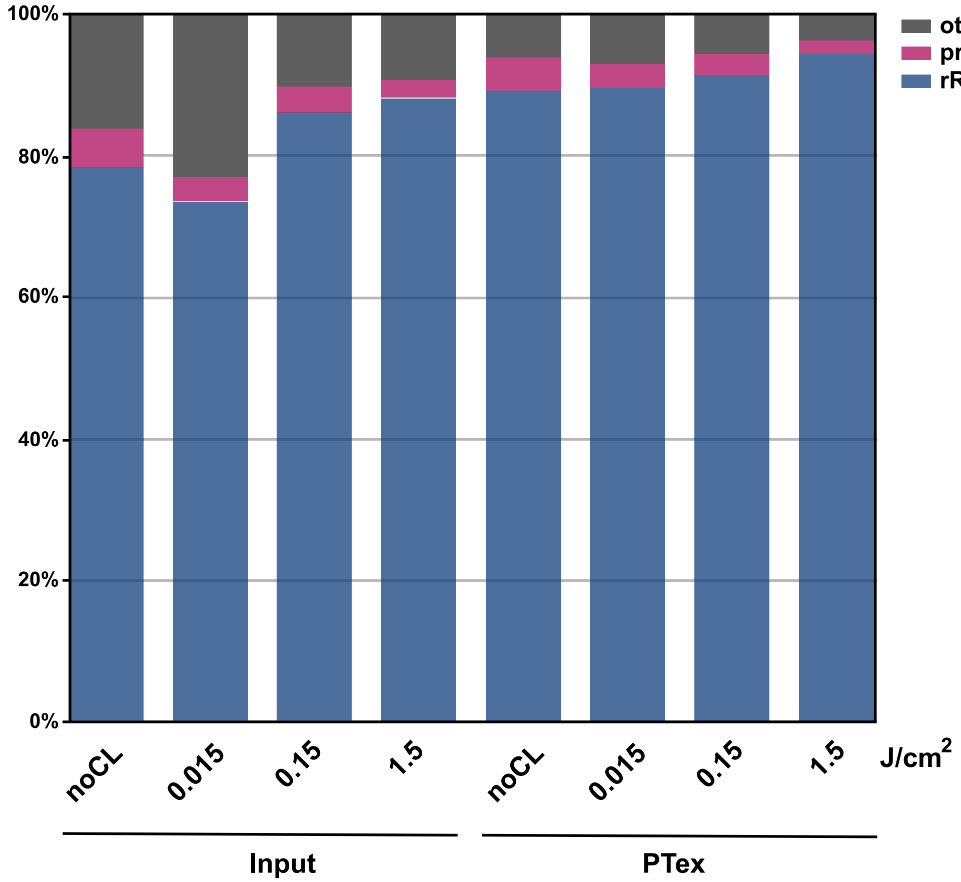


Figure S21: RNA classes of HEK293 total cellular RNA (input) and after PTex purification as analysed by RNA-Seq. Each conditions was sequenced in triplicates (n=3 except for input $0.015 J/cm^2$ (n=2)), shown are mean values. Please note that we omitted ribosomal RNA (rRNA) depletion during the library preparation. PTex-purified RNA largely resembles the cellular distribution of RNA classes, indicating that PTex is not biased for a particular RNA class. We consider all RNA to be interacting with proteins *in vivo*, explaining why PTex-purified RNPs display the same RNA class distribution compared to whole cells. Annotation data are available on Supplementary Table S5.

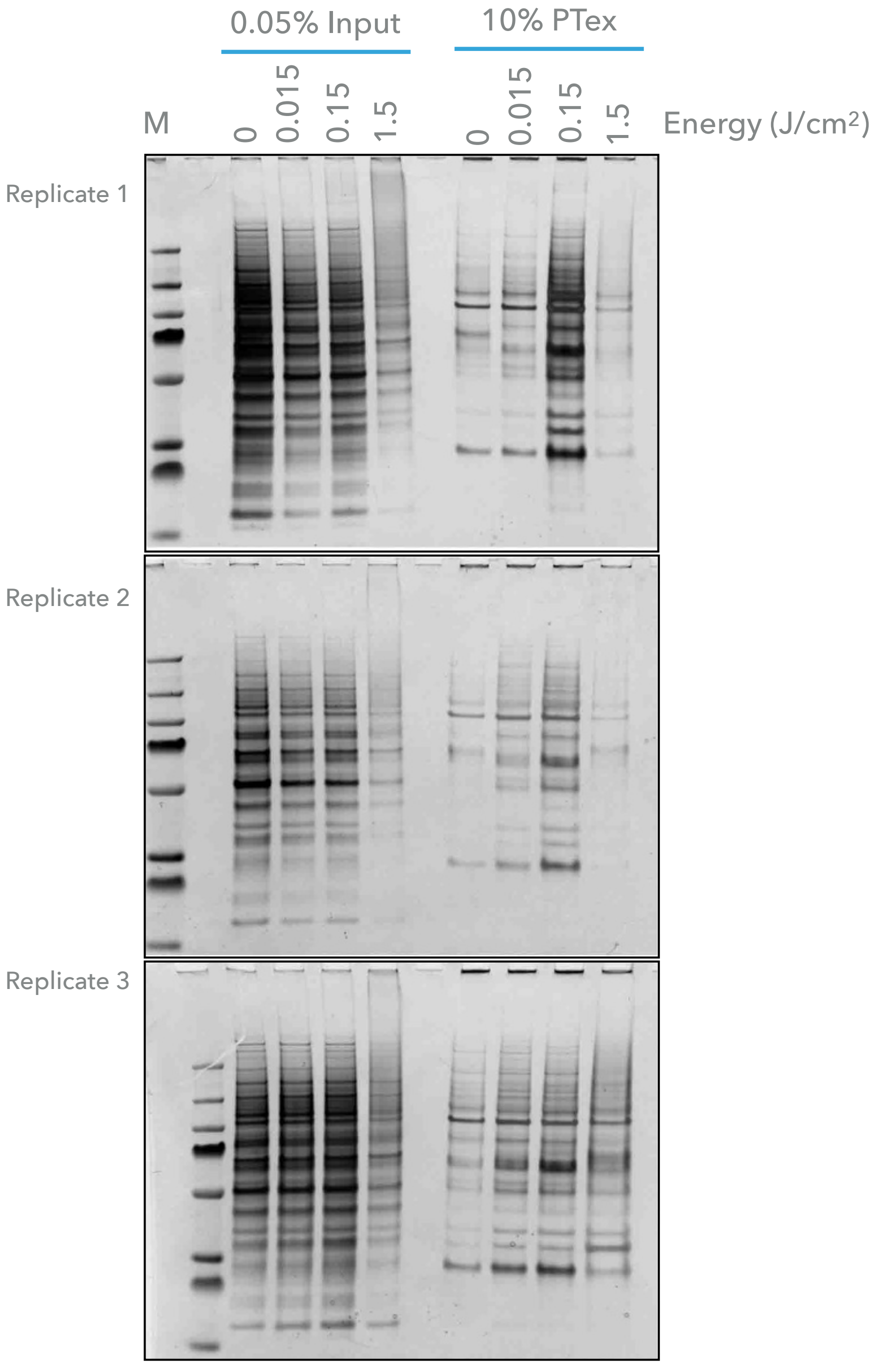


Figure S22: Silver staining of PTex from HEK293 cells during PTex sample preparation for MS (after ethanol precipitation): PTex pellets were solubilised in 2 M Urea and treated with Benzonase, followed by Wessel-Flüggel precipitation, solubilisation in 8 M urea, and digested with DTT, Iodoacetamide, LysC and Trypsin. StageTip desalting was applied.

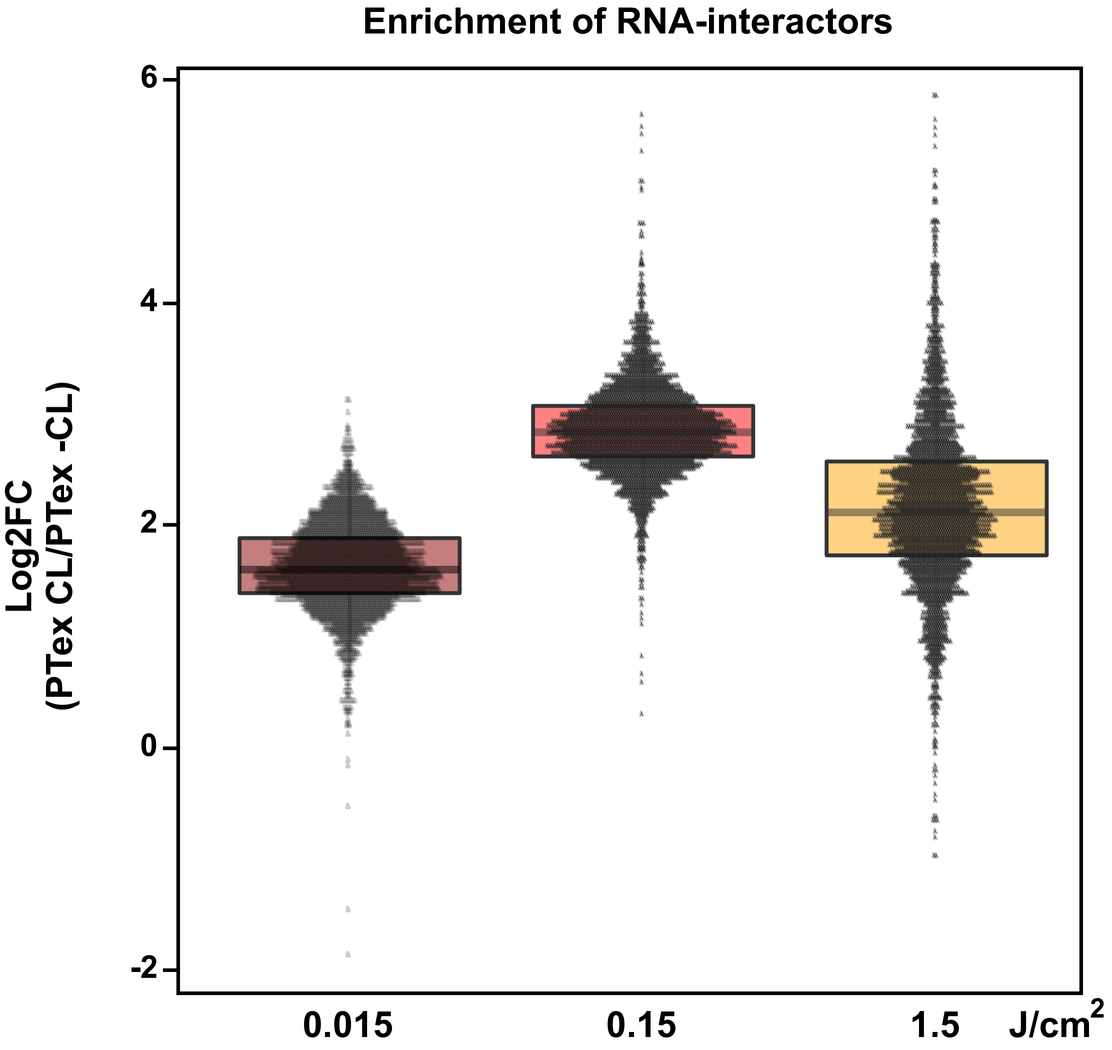


Figure S23: UV intensity vs. PTex enrichment from RNA-associated proteins in HEK293 cells.

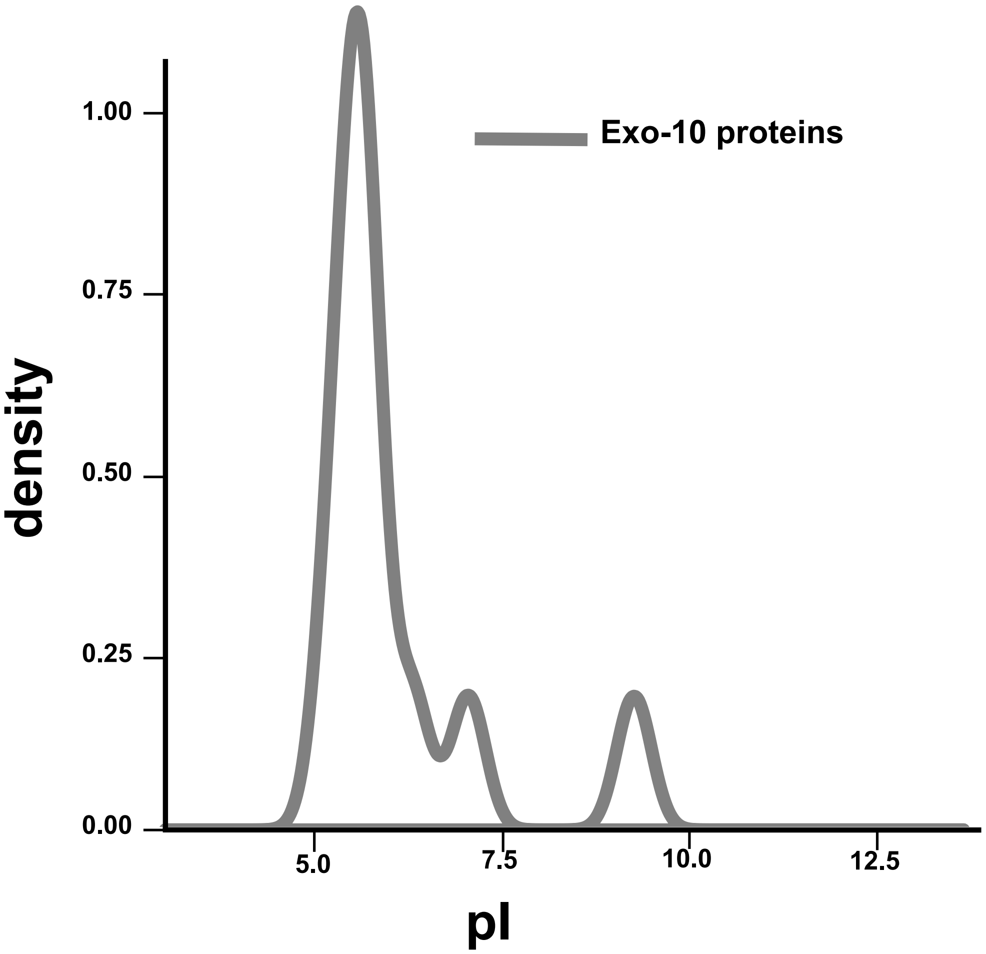


Figure S24: Isoelectric points of human exosome core proteins (Fig. 6).

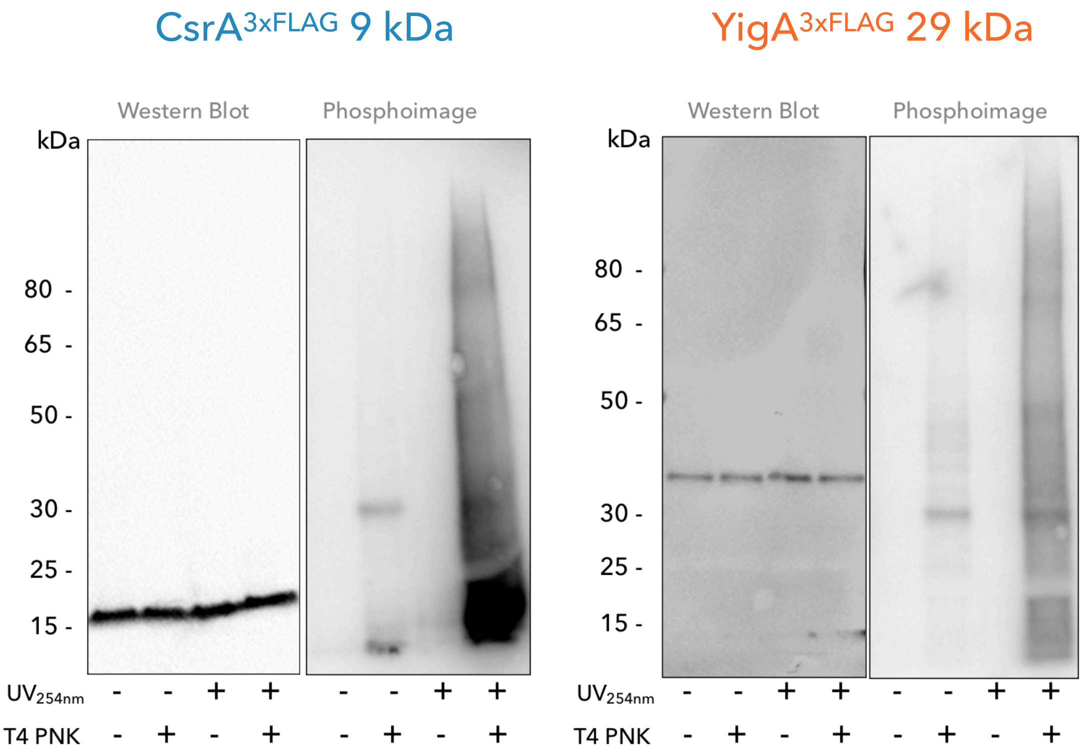


Figure S25: Immunoprecipitation and PNK assay of bacterial RBP CsrA and non-RBP YigA (Fig. 7d).

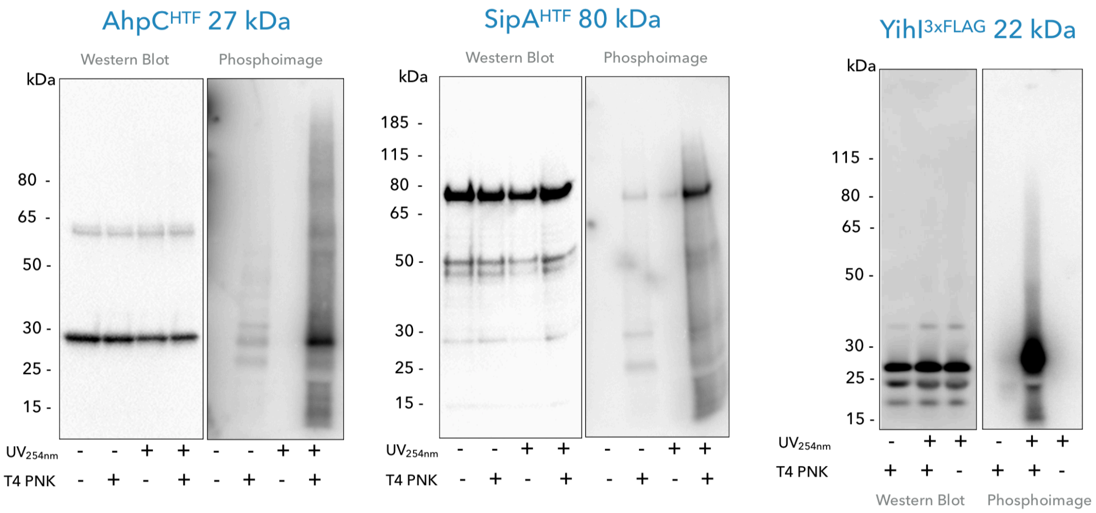


Figure S26: Immunoprecipitation and PNK assay of bacterial RBP candidates AhpC, SipA and YihI (Fig. 7d).

### Supplementary Methods

#### Purification of Crosslinked RNA-Protein Complexes by Phenol-Toluol Extraction (PTex) - Step by step protocol

### Purification of Crosslinked RNA-Protein Complexes by Phenol-Toluol Extraction (PTex)

### Protocol

#### Cells and reagents

- Selected cell line.
- Culture medium according to experiment.
- DPBS (Gibco).
- Solution D (modified from Chomcznski y Sacchi 2006):

    - 5.85 M guanidine isothiocyanate.
    - 31.1 mM sodium citrate.
    - 25.6 mM N-lauryosyl-sarcosine
    - 1% 2-mercaptoethanol.

- Phenol (Roti-Phenol, Roth 0038).
- Toluol (Th.Geyer, 752.1000).
- 1,3-Bromochloropropane (Merck, 8.01627.0250).
- Ethanol absolute.
- Distilled, sterile water.

#### Equipment and Materials

- Incubator (shaker)
- Cross-linking device: 254 nm bulbs (CL-1000, Ultra-Violet Products Ltd.).
- Refrigerated bench-top centrifuge for 2 ml microtubes.
- Refrigerated bench-top centrifuge with rotor for 5 ml tubes.
- ThermoMixer (Eppendorf).
- Microtubes with safe cap (2 ml, 5 ml)
- 1 ml syringes.
- Blunt needles 21G.
- Rubber cells-scrapper.

### Cell culture and crosslinking

Culture cells at 80% confluence in DMEM with D-glucose (10% FCS, 1% P/S). Wash cells in monolayer with 10 ml cold DPBS per dish. Remove the PBS and place the culture dishes without their lids on a cold try inside the cross-linker device. Irradiate with 0.15 - 0.25 J/cm$^2$ at 254-nm UV light. Add 500 µl of DPBS to each dish and scrape the cells with a rubber; place the –CL and +CL pools separately in a 15 mL falcon tube. Take a sample of each for cell counting. Prepare aliquots of 4-8x10$^6$ cells/tube* in 2 ml sterile micro-tubes (500 xg, 3 min, 4 °C). Remove and discard the supernatant, store at -20°C for use within a week, or -80°C for longer.

### PTex: Phenol-Toluol extraction in three steps

**TIMING: ~40 min for extraction / 1h ethanol for precipitation and centrifuging**
**PERFORM ALL STEPS UNDER THE HOOD. USE SAFETY GEAR!**

### STEP ONE

This step must be carried out under physiological conditions. Use of detergents, high concentrations of salts or other denaturing conditions affect the correct behaviour of the complexes during the extraction.

Pre-cool the centrifuge at 4°C. Prepare sets of three 2ml tubes (with safety cap) and one 5 ml tube per sample, label accordingly with an ethanol-resistant marker.

- To one set of 2 ml tubes add 300 uL phenol, 300 µl toluol and 200 µl bromo-chloro-propane (BCP). The second set of tubes will contain 300 µl of solution D, reserve. Add 4.5 ml of Ethanol absolute to 5 ml tubes, reserve. Use the third set of tubes for input control.

- Resuspend –CL and +CL cell pellets in 1 ml DPBS, take 600 µl* and mix with the organic solution during 1 min at room temperature on ThermoMixer at 2.000 r.p.m. From the remaining volume freeze aliquots to be used for gel electrophoresis, DNA or RNA extractions as whole cell input controls.

* In our hands, extraction of more than 8x10$^6$ HEK293 cells per tube caused saturation of the system.

- Centrifuge at 20.000xg, 4° C, 3 min. You will obtain three visible phases: the upper aqueous-phase (aq1), a middle membraneous-like inter-phase (int1), and the organic phase (org1).

- Carefully remove as much aq1 as possible avoiding contact with the int1. Place the aq1 in the corresponding tube with the solution D, mix pipetting up and down (take the same volume of aq1 in all cases). CRITICAL: do not to contaminate the aq1 with the int1. Help yourself with a syringe, blunt needle (21G).

### STEP TWO

- Add 600 µl phenol and 200 µl BCP to the aq1-solutionD tube; mix during 1 min at room temperature, 2.000 r.p.m. (ThermoMixer), and centrifuge as before.

- Carefully with the help of a fresh syringe remove the upper three quarters of the resulting aqueous-phase (aq2) and the lower three quarters of the organic-phase (org2), approximately 600 µl each (keep the same ration among samples). CRITICAL: a whitish-homogeneous layer adhered to the wall of the tube in the aqueous/interphase area might appear, do not disturb or discard it!.

- Keep the resulting interphase in the same tube and proceed with the final step.

### STEP THREE

- Mix the int2 with 400 µl of water and 200 µl of ethanol absolute, shake slightly. Then, add 400 µl phenol and 200 µl BCP, mix and centrifuge as before.

- Remove 3/4th of the aqueous- and organic- phases using fresh syringes. Transfer the remaining inter-phase 3 (200-400 µl) into the 5 ml tubes previously prepared for ethanol precipitation. Cool-down at -20/-80°C during 30 min minimum. PAUSE POINT.

- Precipitate the complexes by centrifuging at 20.000xg, 30min, 4°C. Carefully decant the supernatant and let pellets dry under the hood for no more than 10 min. Resuspend in 30-50 µl of distilled water, Laemmli buffer, or the respective solution according to downstream applications.
